# Altered Hormone and Bioactive Lipid Plasma Profile in Rodent Models of Polycystic Ovarian Syndrome Revealed by Targeted Mass Spectrometry

**DOI:** 10.1101/2024.10.14.618228

**Authors:** Hannah C Scott, Martin Philpott, Georgina Berridge, Zhanru Yu, Darragh P O’Brien, Christian Becker, Cecilia Lindgren, Udo Oppermann, Jörg Müller, Martin Fritsch, Adán Pinto-Fernández, Benedikt M Kessler

**Author notes:** Caeruleus Genomics, Bioescalator Innovation Building, Old Road Campus, Roosevelt Drive, Oxford, OX3 7FZ (Martin Philpott). **Correspondence** (H.C.S) and (B.M.K).

## Abstract

Polycystic ovarian syndrome (PCOS) symptoms include excessive body or facial hair, irregular periods, reduced fertility, and reoccurring pregnancy loss. Hyperandrogenism and chronic inflammation are hallmarks of PCOS, which is diagnosed by analysing steroid hormones in the blood. Studies suggest that bioactive lipids are contributing to chronic inflammation. To research PCOS, animal models, such as letrozole- and dihydrotestosterone-treated rats, are used. They display similar ovarian and metabolic characteristics, although plasma lipid profiles have not been determined. Therefore, in order to validate the use of these models for PCOS, we have optimised a mass spectrometry based targeted lipidomics workflow which increases the sensitivity of measuring these lipids in rat plasma. Our analysis shows that letrozole caused a significant elevation of 5α-androstene-3,17-dione and testosterone. Dihydrotestosterone treatment resulted in increased dehydroepiandrosterone-sulphate and allopregnanolone but a reduction in testosterone, progesterone, pregnenolone, and D-sphingosine. In both models 25-hydroxycholesterol and leukotriene C_4_ were significantly diminished and 4-cholesten-3-one was significantly increased, and these particular metabolites are not known to be changed in human PCOS. These results suggest that the plasma lipids of these rodent models have some changes akin to human PCOS and other changes that are not.

## Introduction

Polycystic Ovarian Syndrome (PCOS) consists of a multifactorial cluster of diseases; the dysregulation of hormone homeostasis underlies reproductive, metabolomic and cardiovascular complications. PCOS is diagnosed based on the presentation of physiological traits, including disturbances in ovulation including oligoovulation or anovulation [1], hyperandrogenism [2], hyperinsulinemia [3], insulin resistance [4], hirsutism [5], acne [6], androgenic alopecia [7], fertility problems, chronic inflammation [8] and polycystic ovaries. PCOS is the most common cause of infrequent or absent periods in women. Preliminary diagnosis of the condition requires steroid hormone analysis to confirm hyperandrogenism [9]. Historically, immunoassays have been used, although more recently, mass spectrometry (MS)-based methods have proven to be more reliable in the identification of PCOS patients with hyperandrogenism [10].

MS methods for steroid and steroid hormone analysis have been developed [11, 12], with targeted approaches such as dynamic multiple-reaction monitoring (dMRM) being able to improve accurate measurement of endogenous species [13]. In particular, MS-based profiling has been performed across many different sample types including serum, plasma, ovarian venous blood, follicular fluid, and various tissues derived from pre-/post-menopausal women, plus endometriosis and PCOS patients [14,15,16]. Measurements indicate that PCOS patients have elevated levels of specific steroid hormones [17] including testosterone [18], 4-androstene-3,17-dione (A4) [19], dehydroepiandrosterone-sulphate (DHEA-S) [20], 5α-androstane-3,17-dione (Aα5) [19], aldosterone [21], cortisone [22], 11-deoxycortisol [23], corticosterone [24], 17α-hydroxyprogesterone [25], 17α-hydroxypregnenolone [26], estriol [27], estrone [28], pregnenolone [24] and dehydroepiandrosterone (DHEA) [29]. In addition, PCOS patients have lower amounts of progesterone [30]. Changes in serum bioactive lipids, in particular Omega-fatty acids (Ω-FAs) upstream of arachidonic acid (AA), eicosanoids downstream of AA [31] and sphingolipids [32] have also been implicated to be aberrant in PCOS patients [33].

Letrozole (LET) is a non-steroidal aromatase inhibitor that interferes with the action of the enzyme CYP19A [34]. CYP19A catalyses the metabolism of testosterone and A4, plus their metabolites to estradiol and estrone respectively. LET is used as an infertility treatment in PCOS patients by inducing ovulation [35]. However, LET is also able to induce PCOS in rats, mimicking human PCOS physiology [36], therefore is used as a model for human PCOS in research. 5α-dihydrotestosterone (DHT) treated rats also display human traits of PCOS pathophysiology and can be used as a suitable alternative model of this disease [37].

Here, we have optimised and applied a targeted analysis for the identification of a selected panel of steroid, steroid hormones and bioactive lipids in the plasma of rats, whereby PCOS is induced by either LET or DHT treatment. Ultra-pressure liquid chromatography coupled to triple quadrupole mass spectrometry (UPLC-MS) using dMRM, was optimised for ultra-sensitivity of detection and for minimising matrix interference. We highlight the challenges of profiling these lipids in complex matrices such as plasma, compare the lipid profiles of these two rat models of PCOS, discuss the differences and similarities to the lipid profiles observed in human PCOS patients and outline potential molecular mechanisms that contribute to the pathophysiology of this syndrome.

## Materials and Methods

### Figures

The **Graphical Abstract, Figures 1B, 3A, and 3B** were created using BioRender.com and **Figure S3** and **Figure S4B** chemical structures were made with ChemDraw version 22.2.0.3300, PerkinElmer Informatics, Inc.

### Reagents

Female Wister Han IGS rats were ordered from Charles River; strain code 273. Rodent diet with 45% Fat [D12451] was purchased from Ssniff. Lipid standards 17α-hydroxypregnenolone [H-105-1ML], DHEA [D-063-1ML], pregnenolone [P-104-1ML], 17α-hydroxyprogesterone [H-105-1ML] and 11-deoxycortisol [D-061-1ML] were purchased from Sigma-Aldrich. LC-MS grade water [115333] and methanol [106035], aldosterone [A9477], progesterone [P0130], Aα5 [A7755-100MG], cortisone [C2755-5G], corticosterone [27840], 5α-dihydrotestosterone [D-073-1ML], testosterone [86500-5G], DHEA-S [700086P-10MG], D-sphingosine (d18:1) [860490P-10MG], leukotriene C_4_ (LTC_4_) [700342E-100UG] and estrone [E9750-5G] were available from Merck. Allopregnanolone [CAY16930-1 mg] was acquired from Cambridge Bioscience Limited. Ammonium acetate [10365260] and isopropanol [15686670] were from Fisher Scientific. Methyl tert-butyl ether (MTBE) was acquired from Acros organics [3787 20010] and acetonitrile was from Honeywell Riedel-de Haën [348512.5L].

Estradiol, Estriol, 25-hydroxycholesterol, dihydrocholesterol and 4-cholesten-3-one were kindly gifted by Professor Udo Oppermann at the Botnar Research Centre (University of Oxford).

### Preparation of steroid and lipid standard solutions

The standard mixture was composed of LTC_4_, estrone, estradiol, estriol, aldosterone, 25-hydroxycholesterol, Aα5, cortisone, testosterone, DHEA, DHEA-S, 17α-hydroxyprogesterone, 11-deoxycortisol, progesterone, dihydrocholesterol, 17α-hydroxypregnenolone, pregnenolone, D-sphingosine, γ-linolenic acid (γ-LA), 5α-dihydrotestosterone, allopregnenolone and 4-cholesten-3-one.

Individual lipid standards were prepared in 100% MeOH, then pooled and diluted to a final concentration of 50 nmol/L.

### In vivo models

Animal experiments with rats were performed as described previously [37, 38]. In brief, female rats were purchased at an age of 2 weeks, with their foster mothers, from Charles River Laboratories Inc, Sulzbach, Germany. Rats were raised until 21 days of age and then used for experiments. Rats were randomly divided into three experimental groups; control (n = 6), DHT (n = 3), and LET (n = 3) and implanted subcutaneously with continuous-release implants (Bayer AG, Berlin, Germany) containing either poly-lactic-glycolic acid (PLGA) matrix only (control), 18 mg LET in PLGA at a daily dose of 200 µg, or 11 mg DHT in PLGA at a daily dose of 80 µg. Rats were kept on a high fat diet from Sniff until the study was concluded. When the rats were 49 days of age blood samples were taken to isolate plasma, which was prepared by lithium heparin protocol. Plasma was kept at -80°C until lipid extraction and MS analysis. Protocols of accepted standards of animal care were approved by the Landesamt für Gesundheit und Soziales (LaGeSo, Berlin, Germany) under the animal allowance number A 0077/17.

### Lipid Extraction

A small amount of coagulation (a clot) was seen in plasma samples; LET-C biological replicates 1, 2 and 3 and LET biological replicates 1, 2 and 3. Plasma (25 µL) material was pipetted as to avoid the coagulation clot and processed further for MS analysis.

Collected plasma material was thawed on ice (4°C), lipids were then extracted with the addition of 100 µL of 50% methanol in water, followed by the same volume of MTBE, vortexed and then mixed at 4°C for 20 min with gentle rotation. The sample was then centrifuged at 17 x G at 4°C for 5 min. The upper organic layer was removed and dried by vacuum centrifugation (speedvac). Samples were stored at -20°C until LC-MS analysis. Before analysis, the lipid extracts were reconstituted into 40 µL of methanol.

All samples were pooled by combining a small portion of each sample to create a “Matrix sample” for determining the matrix effects. Two “Matrix” samples were extracted as described above by extracting from 20 µL of the pooled plasma. To one of the “Matrix” samples the methanol in the extraction protocol was substituted for the standard mixture, see section Materials and Methods, **Preparation of steroid and lipid standard solutions**, to create a “Matrix plus standards” sample.

### LC-MS Method

Levels of LTC_4_, estrone, estradiol, estriol, aldosterone, 25-hydroxycholesterol, Aα5, cortisone, testosterone, DHEA, DHEA-S, 17α-hydroxyprogesterone, 11-deoxycortisol, progesterone, dihydrocholesterol, 17α-hydroxypregnenolone, pregnenolone, D-sphingosine, γ-LA, 5α-dihydrotestosterone, allopregnenolone and 4-cholesten-3-one were analysed. Metabolites were quantified by liquid chromatography mass spectrometry (LC-MS) using an optimised dMRM method on a triple quadrupole mass spectrometer with a JetStream ESI source (Agilent 6495) coupled to a 1290 Agilent LC system.

Lipids were separated on an ACQUITY UPLC BEH C18 column (1.7 µm, 100 × 2.1 mm i.d., Waters). Mobile phase A was 40% acetonitrile with 5mM ammonium acetate and mobile phase B was 90% isopropanol, 10% acetonitrile with 5mM ammonium acetate. The flow rate was set to 0.2 mL min−1 and the sample injection volume was 10 µL. The following gradient (% mobile phase B) was used: 0–2.5 min at 2% B, 2.5–25 min 100% B. A wash with 100% mobile phase B was performed to clean the column before re-equilibration to starting conditions. The autosampler was maintained at 4°C.

The following ESI source parameters were used: gas temp at 280°C, gas flow 14 L/min, nebuliser at 20 psi, sheath gas temp at 250°C, sheath gas flow at 11 L/min, capillary voltage 4000 V, nozzle voltage 1000 V, high-pressure RF at 150 V and low-pressure RF at 90 V. The MS was operated in fast switch mode, changing polarity from positive to negative, depending on the dMRM transitions. The dMRM fragmentor voltage (V) was set at 380 and the cell accelerator voltage at 4. The transitions used in the dMRM analysis are shown in **Table S1**; the LC and MS parameters used are shown in **Table S2.**

### Data Analysis

Initial data processing was performed using Agilent MassHunter Quantitative Analysis software (v. 10). Post-processing was performed in Excel and GraphPad Prism, in which dMRM responses were corrected with a blank subtraction by removing any response in a methanol only injection. Responses were corrected for drift in instrument performance by adjusting to a quality control sample and then modified by the matrix response factor (matrix RF). To determine the matrix response, and to correct for any matrix effects we used our previously published protocol for data analysis [39]. Matrix RF-corrected response can subsequently be used to calculate analyte concentration from a standard curve trendline equation (y = Mx + C). Standard curves are generated by the injection of increasing concentrations of the lipid standard solution, calculating the response at each concentration, and plotting the values against the corresponding concentration.

Statistical analysis of the two sample or unpaired t-test is used to validate difference between no treatment (Control) and treated with LET or DHT. A p-value less than 0.05 (*) indicates that the results have highly significant differences. If there is no significance difference it is unlabelled.

### Method Validation

The lower limit of detection (LLOD) was calculated as a signal-to-noise (S/N) ratio of >3; the lower limit of quantification (LLOQ) was a S/N ratio of >10. Intraday precision was calculated using three replicates of two concentrations over the course of one day and the results are reported as coefficient of variation (CV in %) between replicates of one concentration. To monitor instrument performance over time and check that analysis that spanned inter-day were consistent a quality control sample, which contained 3.5 fmol/L on-column (o.c.) of testosterone was routinely injected. If the total sample analysis time was over multiple days, the samples were briefly vortexed at the start of each day to avoid precipitation. Variation in biological and technical replicates was investigated and results are shown in **Figures S1** and **S2**, **Tables S3** and **S4**.

## Results

Lipid standards were selected based on availability and to cover metabolites representing steroids, steroid hormone biosynthesis and degradation, eicosanoid metabolism and sphingolipid metabolism (**Figure 1A**). Our panel included lipids from the cholesterol, progestin, mineralocorticoid, glucocorticoid, androgen, estrogen, fatty acid, eicosanoid and sphingolipid families (22 in total, **Figure S3**). We developed a lipidomics workflow for analysis of these lipids at low concentrations and for minimal matrix interference in rat plasma (**Figure 1B**). Chromatographic separation of each of the lipid standards were optimised and their detection levels maximised in ESI ionisation MS. Tandem MS (MS^2^) spectra were generated to determine diagnostic fragment ions, as exemplified for testosterone **(Figure S4A**). The protonated mass of testosterone was determined at mass-to-charge (*m/z)* 289, the sodium adduct is at *m/z* 311 and a protonated dimer at *m/z* 577.1, respectively. Endogenous sodium contribution may be from the plasma, although this is removed by the lipid extraction process as the lower aqueous fraction is discarded. We used this information to cross reference our spectra with those found in public databases, such as Human Metabolome Database and LipidMaps, to corroborate identification of the correct lipid. We established the retention time of the lipids by MS^2^ to determine a retention time window for the application of specific dMRM settings (**Figures S4B and S4C).** MS-based quantitation of the lipid standard mixture (see section Materials and Methods, **Preparation of steroid and lipid standard solutions**) was established by a calibration curve of the quantification ion, demonstrating linearity over ∼2 orders of magnitude, reaching 1 fmol/L o.c. (**Figure S4D)**. Representative dMRM chromatograms for both quantification and qualification ions for all the lipids studied were obtained by optimising parameters for each individually (**Figure 1A**, **Table S2)**. This resulted in a compendium of dMRM settings applied for quantification of the panel of lipids profiled in this study **(Table S1**). The response curve equations and correlation coefficients varied from 0.917 for 5α-dihydrotestosterone to 0.999 for 11-deoxycortisol, Aα5, 17α-hydroxyprogesterone, progesterone, estrone, and testosterone, reflecting accurate quantitation over the defined concentration range **(Table S5**). Once optimal MS parameters were determined, we subsequently validated the reproducibility and robustness of our methodology by performing an intra-day variation analysis (**Figure S5**). Lipid standards were combined and analysed twice at 100 fmol o.c. The variation, in percentage, from the first analysis was calculated for the second independent analysis. The observed variations for all the lipids were lower than 5% for most lipids, highlighting the reproducibility of the method. To detect the potential presence of isomers, we performed a transition ion ratio (TIR) analysis **(Table S6**). Testosterone, Aα5 and DHEA are isomers having the same molecular mass, therefore chromatographic separation is essential to provide accurate quantitative results. Testosterone and DHEA are chromatographically separated, removing the need for a TIR analysis in this case, but testosterone and Aα5 are only slightly separated. Therefore, the instability of the ratio observed in the TIR of testosterone could be due to Aα5 interference. However, this issue is mitigated by using different quantitation ion transitions for testosterone and Aα5.

**Figure 1A:**
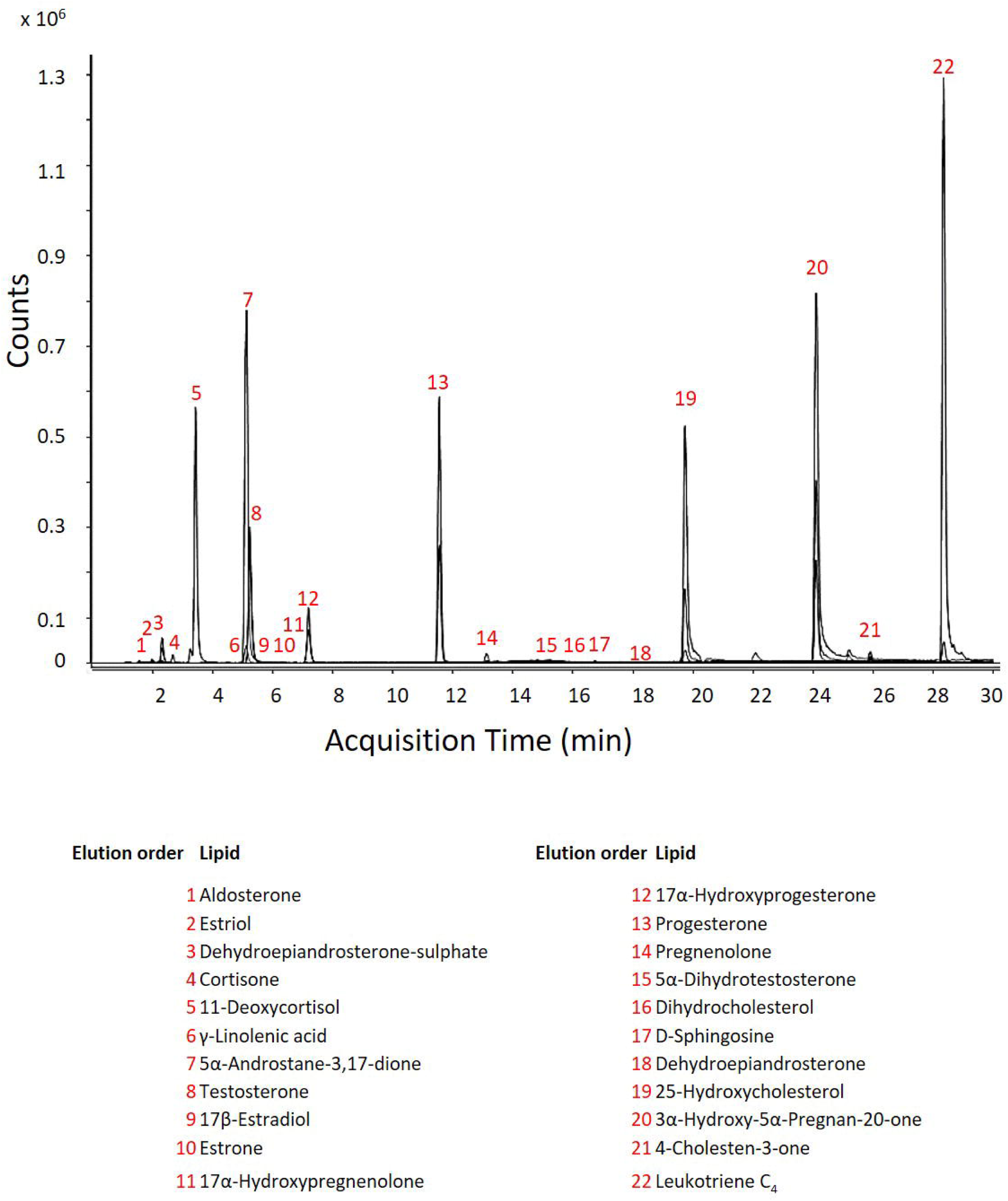
Optimised lipid chromatography. The dMRM chromatograms for both quantitation and qualification ions for all the lipids analysed with this methodology. The lipids are analysed at the same on-column concentration of 300 fmol/L, the difference in counts for each lipid highlights the differences in ionisation efficiencies of the lipids (molecular structures of analysed lipids are in **Supplementary Figure 3**).

**Figure 1B:**
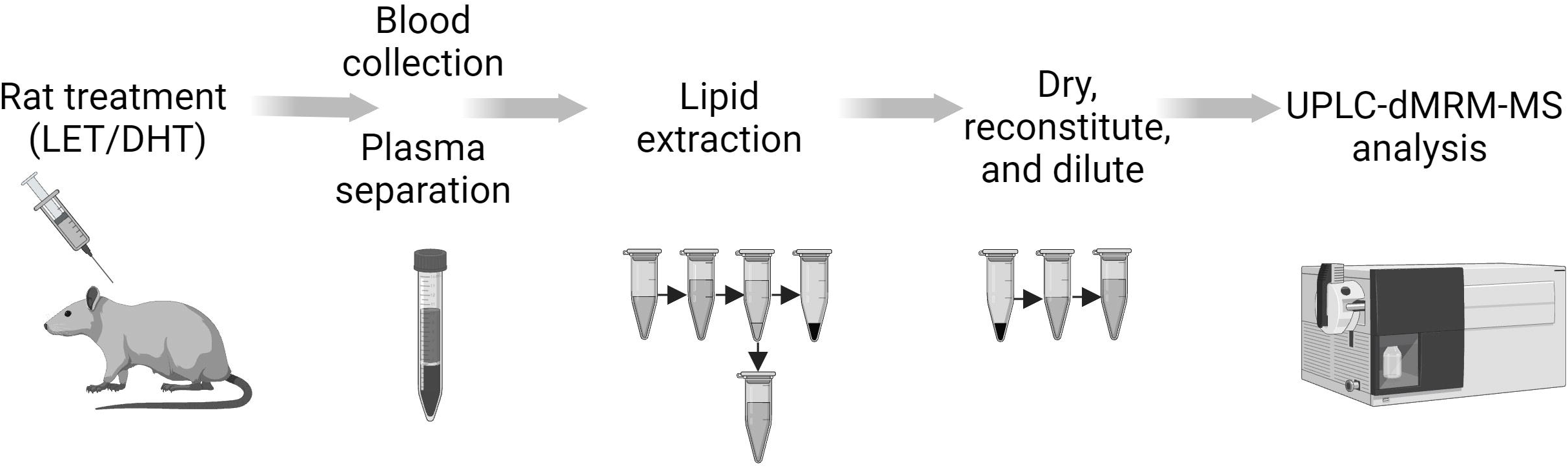
Optimised experimental workflow for the analysis of steroids, steroid hormones and bioactive lipids in rat plasma. Analyses were conducted using an ultra-high-performance liquid chromatography-dMRM-MS (UPLC-dMRM-MS) analysis. Rat are treated with either Letrozole (LET) or Dihydrotestosterone (DHT) to induce PCOS-like physiology. At age 49 days, blood was taken from the rats and plasma was obtained by Lithium-Heparin separation. Lipid extraction from plasma was performed with Methanol:Methyl-tert-butyl-ether (50:50, v:v). Lipid extracts were subsequently dried and reconstituted into Methanol prior to LC-MS analysis.

Having optimised analytical parameters, we set out to discover potential alterations of lipid profiles observed between individual rats for each treatment condition; LET, Control for LET (LET-C), DHT and Control for DHT (DHT-C), (**Figure S1**). The CV for biological variation for the profiled lipids ranged from 1.06 % for allopregnanolone (3α-hydroxy-5α-pregnan-20-one, AP) measurement in LET-C treated rats, to 141.42% for Aα5 measurement in DHT-C rats **(Table S3)**. Trace amounts of Aα5 in DHT-C was responsible for the high variation. It is worthwhile to note that LET-C and LET rat plasma had coagulation, and this may contribute to the biological variation seen (Materials and Methods Section - Lipid Extraction).

Technical triplicate analysis of each biological replicate revealed tighter consistency (**Figure S2)**. The CV for technical variation had a range of 0.02% for progesterone analysis in DHT-C biological replicate 1 to 90.23% for Aα5 analysis in LET-C biological replicate 3 (**Table S4**). Trace amounts of Aα5, were responsible for the high variation in LET-C. Aα5 was not present in DHT biological replicates 1-3 or in DHT-C biological replicates 1 and 2, therefore no technical variation analysis could be performed. The biological condition LET showed an average technical replicate CV of 7.34%, LET-C had 10.44%, DHT had 5.95% and DHT-C showed 7.30%. In conclusion, having established the variation of lipid measurements in rat plasma, we could then assess quantitative changes and their significance upon treatment conditions.

LET treatment of rats resulted in marked changes to the lipids profiled in rat plasma. We observed significant increases in the concentration of Aα5 (+ 135.75 fmol/L, p=<0.0001), testosterone (+ 150.71 fmol/L, p=<0.0001) and 4-cholesten-3-one (+ 72.53 pmol/L, p=0.0063). DHEA-S, 11-deoxycortisol, pregnenolone, D-sphingosine and 3α-AP were also increased in LET treated rat plasma, although these metabolites were not significantly elevated **(Figure 2A**, **2C, 3A, Table S7)**. A significant decrease in the plasma levels of 25-hydroxycholesterol (- 4.21 pmol/L, p=<0.0001), and LTC_4_ (- 2.34 pmol/L, p=0.0027) were observed. Progesterone and γ-LA were also reduced in LET treated rat plasma, although these metabolites were not significantly decreased (**Figure 2A**, **Table S7**).

DHT treatment of rats also resulted in significant changes to the lipids profiled. For instance, we observed significant increases in the concentration of DHEA-S (+ 225 fmol/L, p=0.024), allopregnanolone (+ 24.67 pmol/L, p=0.0031) and 4-cholesten-3-one (+ 450.52 pmol/L, p<0.0001) **(Figure 2B**, **2C, 3B, Table S7)**. 11-deoxycortisol, and γ-LA were also increased in DHT treated rat plasma, although these metabolites were not significantly elevated. Notably, a significant decrease in the plasma levels of testosterone (- 4.55 fmol/L, p=0.0012), progesterone (- 332.34 fmol/L, p=0.0044), pregnenolone (- 12.90 pmol/L, p=0.0018), D-sphingosine (- 617 fmol/L, p=0.047), 25-hydroxycholesterol (- 6.20 pmol/L, p=<0.0001), and LTC_4_ (- 4.32 pmol/L, p=0.0003), were observed. Aα5 was also decreased in DHT treated rat plasma, although this androgen was not significantly reduced. Aside from two data points (2 technical replicates of 1 biological replicate), it is all but absent in both DHT and DHT-C samples (**Figure 2B**, **Table S7**), contributing to the high variation seen in the analysis of this androgen.

**Figure 2A:**
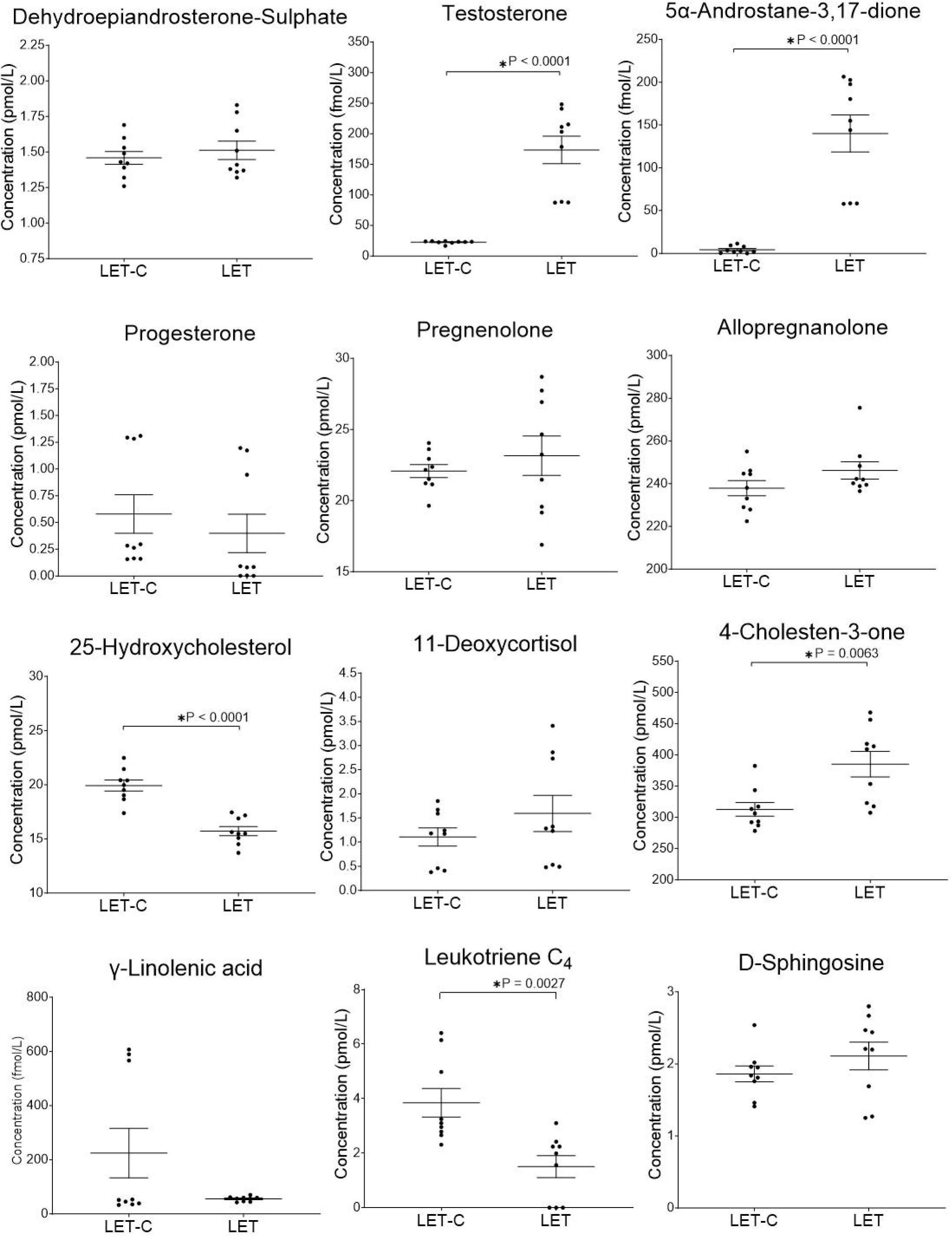
Analysis of lipids in rat plasma after LET treatment. Letrozole (LET) induced PCOS in rats modulates steroid hormone biosynthesis, steroid hormone degradation, eicosanoid and sphingolipid pathways. The Letrozole control (LET-C) is on the left and the Letrozole (LET) treated is on the right. Black dots are each data point (technical triplicate of biological triplicates, n=9). The statistical analysis is the two sample (or unpaired) *t*-test. A *p*-value less than 0.05 (*) suggests that the results have highly significant differences. If there is no significant difference, it is not labelled. **Supplementary Table 7** contains the values for mean, standard deviation (SD) and population number (n=).

**Figure 2B:**
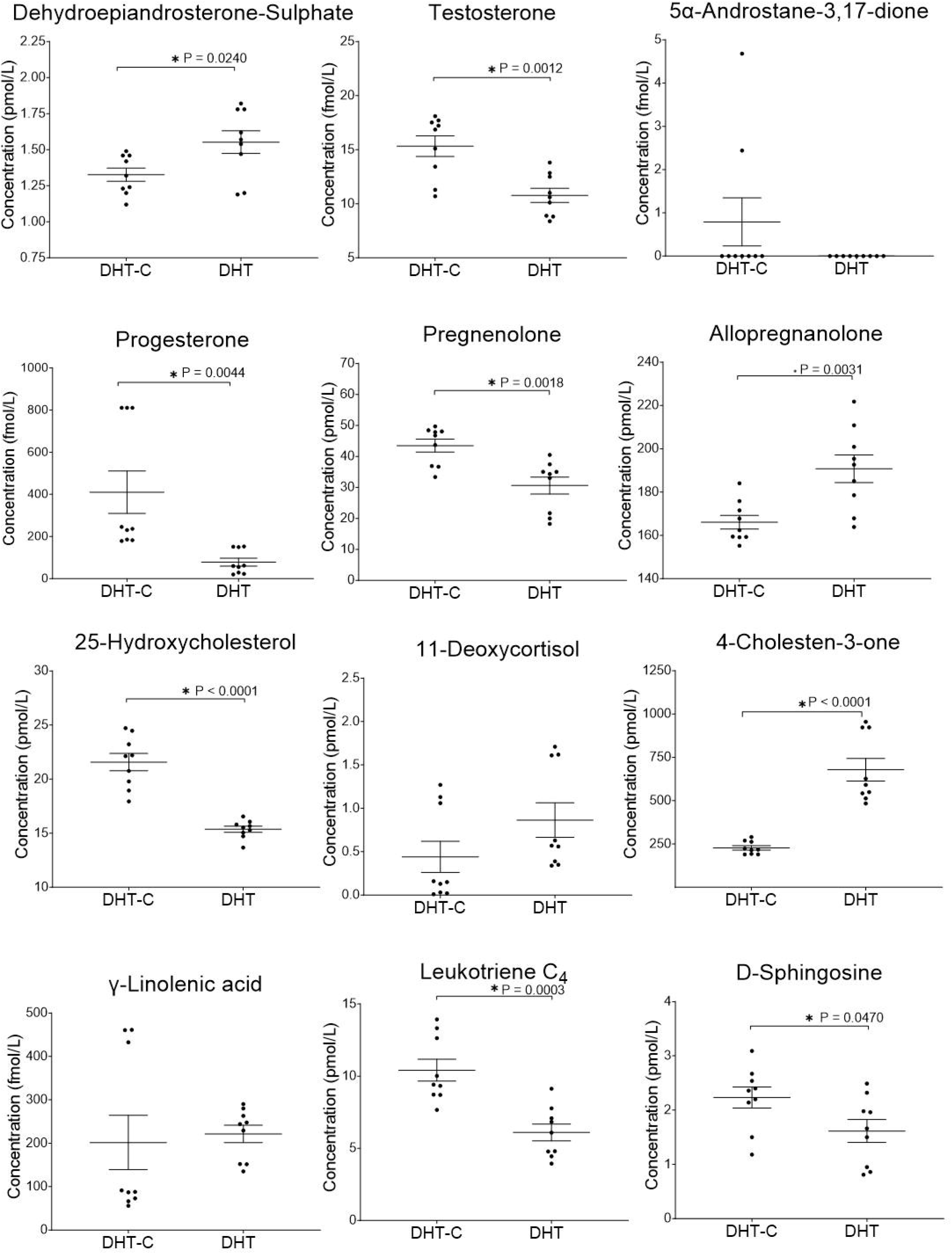
Analysis of lipids in rat plasma after DHT treatment. Dihydrotestosterone (DHT) induced PCOS in rats modulates steroid hormone biosynthesis, steroid hormone degradation, eicosanoid and sphingolipid pathways. Graphs represent the mean (line) with the standard error of the mean (error bars). The DHT control is on the left and the DHT treated is on the right. Black dots are each data point (technical triplicate of biological triplicates, n=9). The statistical analysis is the two sample (or unpaired) *t*-test. A *p*-value less than 0.05 (*) suggests that the results have highly significant differences. If there is no significance difference, it is not labelled. **Supplementary Table 7** contains the values for mean, standard deviation (SD) and population number (n=).

**Figure 2C:**
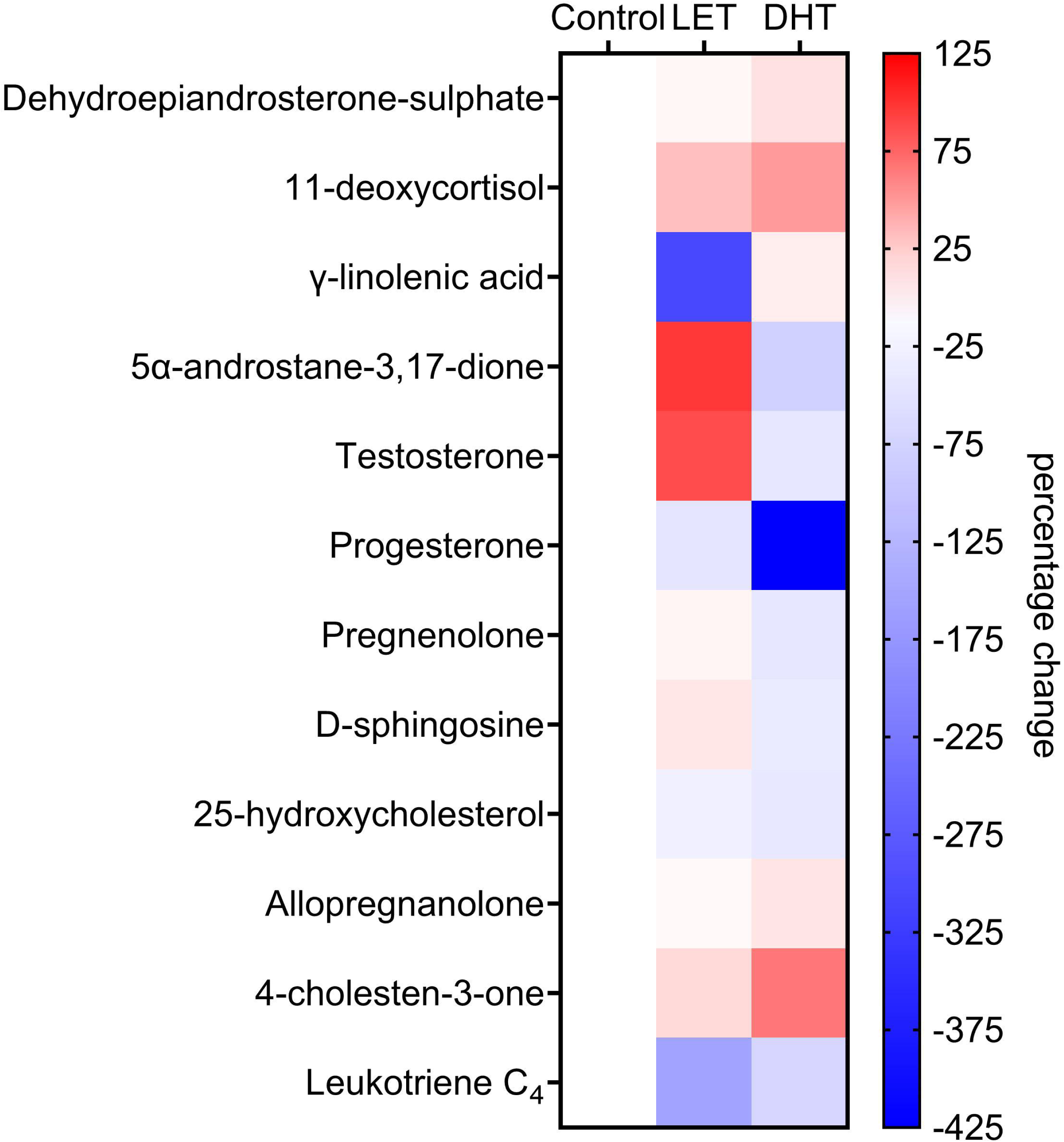
Analysis of lipids in rat plasma after Letrozole (LET) or Dihydrotestosterone (DHT) treatment. Mean percentage change of lipids in rat plasma, not-treated control (Control) and treated with LET or DHT. Heat maps display normalised lipid abundances relative to the mean concentration of the respective non-treated control samples (Control), which is represented by the white boxes. For LET or DHT treatment, blue results are for a decrease in concentration, in percentage, white is for no change and red is for an increase in concentration in percentage.

**Figure 3A:**
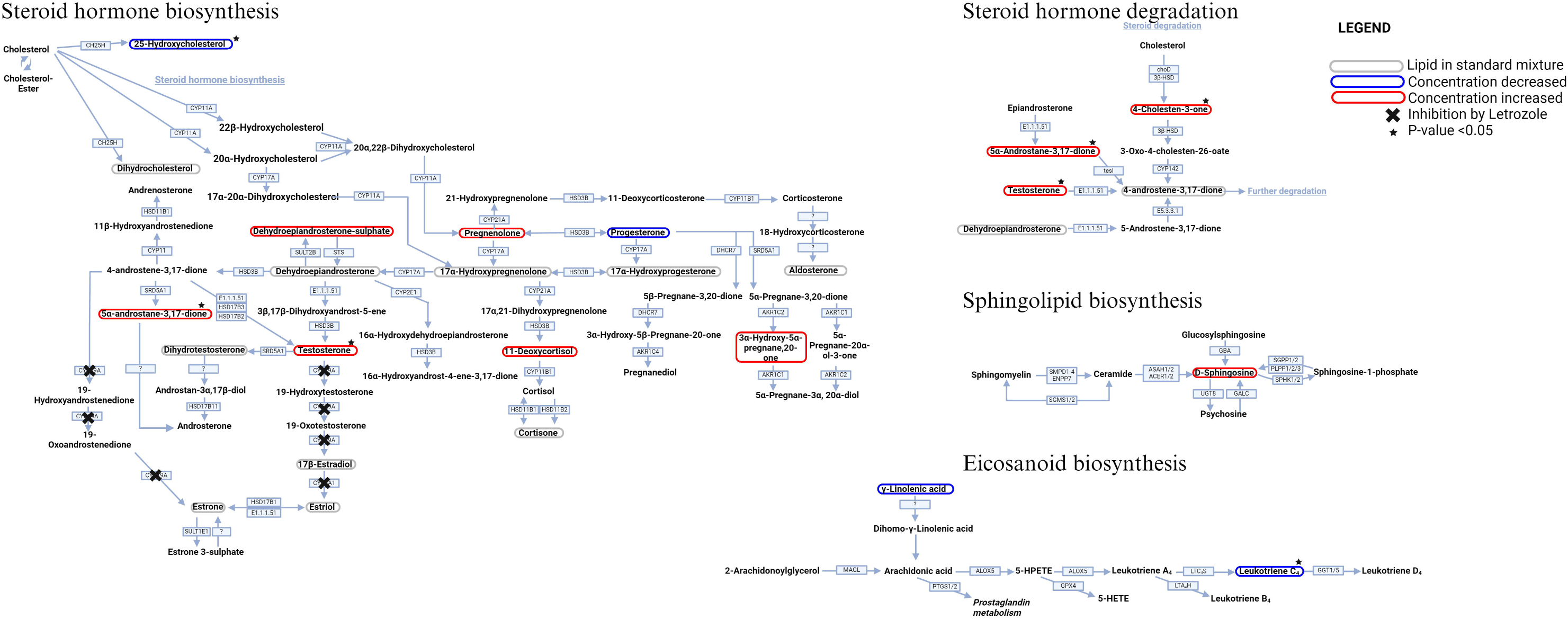
Letrozole (LET) treatment pathway analysis. Lipid profile after treatment with LET. Converting enzymes are in boxes and the lipids in this study are circled. Lipids circled in grey have either no change compared to non-treated control rat plasma or were not detected with this methodology; lipids circled in blue have a decreased concentration compared to non-treated control rats; and lipids circled in red have an increased concentration compared to non-treated control rats. Lipids with a significant difference between non-treated control and LET treated (a *p*-value of less than 0.05) are indicated by a black star and inhibition of CYP19A activity by LET with a black cross.

**Figure 3B:**
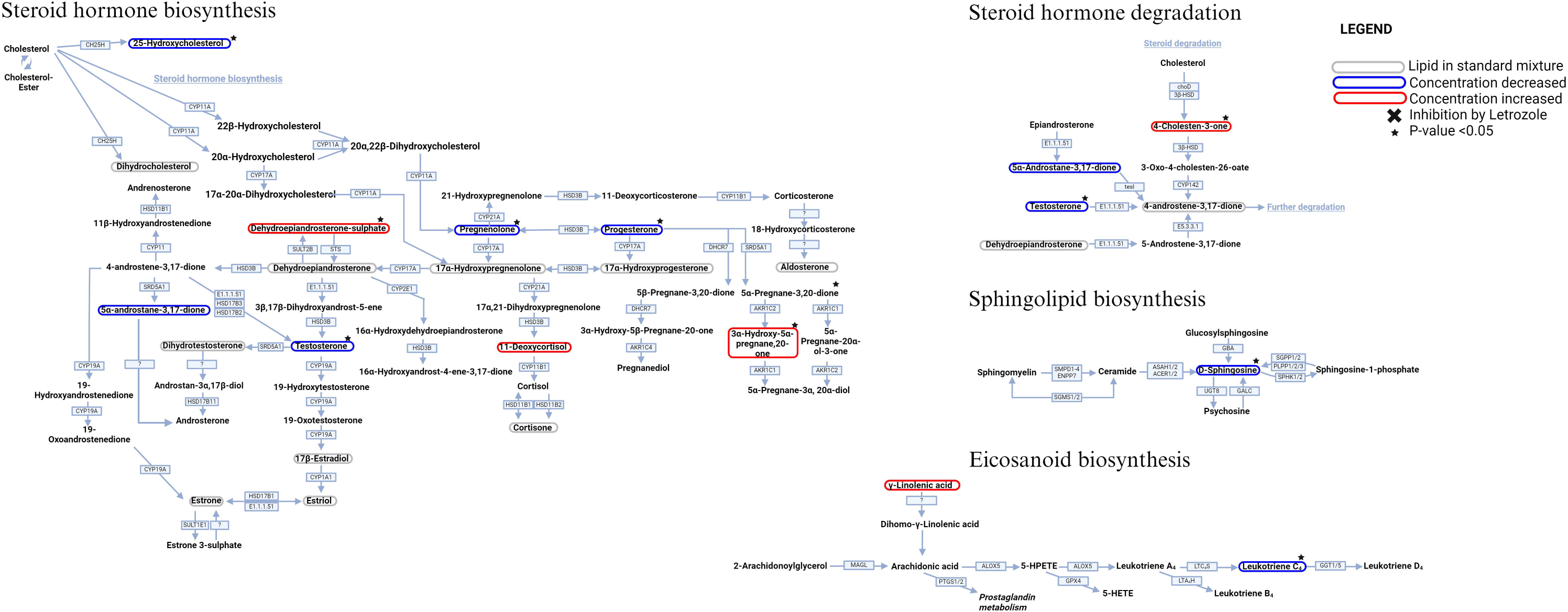
DHT treatment pathway analysis. Lipid profile after treatment with DHT. Converting enzymes are in boxes and the lipids in this study are circled. Lipids circled in grey have either no change compared to non-treated control rat plasma or was not detected with this methodology; lipids circled in blue have a decreased concentration compared to non-treated control rats; and lipids circled in red have an increased concentration compared to non-treated control rats. Lipids with a significant difference between non-treated control and DHT treated (a p-value of less than 0.05) are indicated by a black star.

Taken together, our optimised targeted mass spectrometry method reported considerable changes in rat plasma steroid hormone and bioactive lipids upon LET and DHT treatment.

## Discussion

Here, we investigated steroid hormones and pro-inflammatory lipid mediators in PCOS rat models to understand its underlying pathophysiology. The lipids analysed in the rat models are representative of lipids involved in steroid hormones biosynthesis and inflammation. We choose bioactive lipids with known involvement in inflammation, and immunity to study this phenomenon in these two models.

Lipidomics remains challenging and continues to benefit from improved methodologies in detection, identification and sample preparation using applicable matrices [39]. Alternatively, a surrogate matrix for steroid hormone analysis may be used, e.g. charcoal stripped samples [40]. However, pre-analytical influences may change lipid concentrations or matrices, which is why we prefer to use extensive matrix-dependent normalisations for each analyte within a “true” matrix, and not use a surrogate matrix. We demonstrate its application tailored to specific endogenous lipid detection in research relevant biological samples matrices as well as plasma in the present study.

Stable isotope labelling [41] or derivatization methods [42] may provide increased sensitivity for quantitative analysis of steroid hormones. Isomeric elucidation, such as determining the identification of, and causing the separation of Aα5 and 5β-Androstane-3,17-dione would be possible by using a MS equipped with ion mobility separation [43]. Alternatively, for determining the presence of isomers in biological matrices the use of a Transition Ion Ratios (TIR) analysis is a possibility. However, TIR is based upon fragmentation of standards, and qualification ion fragments for standards are not always present in biological samples. This can be due to falling below the lower limit of detection (LLOD) or lower limit of quantification (LLOQ), matrix effects and other analytical singularities including concentration dependence [44]. Our preference is to use qualification ions solely when the identity of the lipid is ambiguous, however our use of standards for confirmation of identification prevents this. In our analysis, not all qualification ions for every lipid profiled are present in all the samples (**Table S6**). Therefore, we believe that the application of TIR may not provide further advantages, at least for the study we describe here. TIR analysis of progesterone suggests the presence of isomers, however, the potential isomer of progesterone is currently unknown. Our method chromatographically separates the isomer species aldosterone and cortisone (C_21_H_28_O_5_), plus testosterone with DHEA and Aα5 (C_19_H_28_O_2_), allowing for accurate identification. However, for the isomer pairs androstenedione with 5α-dihydrotestosterone (C_19_H_30_O_2_), 11-deoxycorticosterone with 17α-hydroxyprogesterone (C_21_H_30_O_3_, 330.5 g/mol), and pregnenolone with 5α-dihydroprogesterone (316.49 g/mol), the situation is less clear. We did not include the standards for androstenedione, 11-deoxycorticosterone or 5α-dihydroprogesterone within our standard mixture, so are currently unable to know if these lipids are chromatographically resolved. We were also unable to detect 5α-dihydrotestosterone or 17α-hydroxyprogesterone in these samples. We are therefore unable to determine if the responses of 5α-dihydrotestosterone, 17α-hydroxyprogesterone or pregnenolone within our samples are contributed solely by these lipids or also include a contribution from their isomers. Measurement of 17α-hydroxyprogesterone, 17α-hydroxypregnenolone, corticosterone, dihydrocholesterol, DHEA, aldosterone, cortisone, 5α-dihydrotestosterone, estradiol, estrone and estriol were not detected in these samples, indicating either their absence, that they fell below the LLOD, and/or the matrix caused considerable ion suppression. Certain metabolites, such as LTC_4_, may benefit from a more tailored analysis that increases sensitivity and quantitation [39].

Despite natural biological heterogeneity, our results show clear differences in the concentration of Aα5 in LET treated rats as well as differences in DHEA-S, progesterone, pregnenolone, allopregnanolone and D-sphingosine quantities in the plasma of DHT treated rats as compared to controls. Both LET and DHT rats display considerable changes in testosterone, 4-cholesten-3-one, 25-hydroxycholesterol and LTC_4_. For both, LET and DHT treatment in the case of increased metabolite concentrations, this could suggest an increase in the synthesis of these lipids, a reduction in their metabolism or conversion, their cellular excretion maybe reduced.

Allopregnanolone (3α-hydroxy-5α-pregnane-20-one, AP) is a pregnane neurosteroid, which has been implicated in reproductive-related psychiatric disorders, such as post-partum depression [45] and premenstrual disorders [46, 47]. Women with PCOS experience increased rates of depression and anxiety [48]. Neuroactive steroids synthesised in the brain and/or crossing the blood-brain barrier (e.g., allopregnanolone and DHEA-S) may affect the serotonergic and glutamatergic systems, and act as allosteric agonists of the GABA_A_ receptor, affecting sleep, mood and appetite [49]. Both allopregnanolone and DHEA-S have been implicated in the mental health complications and overeating caused by, or associated with PCOS [50, 51]. However, human and animal models of PCOS have mixed and sometimes conflicting results on the contribution of allopregnanolone in PCOS development and pathology [52, 53]. In our analysis, progesterone is decreased in LET and DHT-treated rats, and allopregnanolone levels are elevated. Progesterone levels may be reduced due to conversion to 5α-dihydroprogesterone, and, subsequently, to allopregnanolone. DHEA-S is increased in both, LET-and DHT-treated rats as compared to controls. High allopregnanolone and DHEA-S amounts may be correlated with weight gain, psychiatric complications and associated metabolic syndromes in these animal models.

4-cholesten-3-one is a cholesterol metabolite involved in the steroid degradation pathway. It can reduce breast cancer cell viability, lipogenesis and enhance the expression of Liver X Receptor-dependent cholesterol transporters, including ABCA1 and ABCG1 [54]. 25-hydroxycholesterol, a primary bile acid, can be transported into cells by ABCA1/G1 [55]. Both rat models have high plasma levels of 4-cholesten-3-one and low amounts of 25-hydroxycholesterol, which may suggest the reduction of 25-hydroxycholesterol in the plasma is due to an increased expression of ABCA1/G1 by high 4-cholesten-3-one which has caused the cellular uptake of 25-hydroxycholesterol by ABCA1/G1.

CYP19A inhibition by LET treatment caused an increased concentration of testosterone, as it cannot be converted into downstream metabolites. Conversely, DHT treatment reversed the effect, with a significant decrease in testosterone. This could be because DHT and testosterone are within the same metabolic pathway, and high levels of DHT may cause a negative feedback loop, leading to a decreased production of testosterone, to control DHT levels. The decrease in testosterone observed in DHT treated rats may also be the result of testosterone glucuronidation, by uridine diphosphoglucuronosyltransferase enzymes, which lowers plasma testosterone levels through secretion of testosterone-glucuronidation products [56].

In addition to LET inhibition of CYP19, Aα5 is also capable of inhibiting CYP19 activity. Reduction in CYP19 catalysation reduces estrogen production from androgens. Aα5 has been shown to be elevated in PCOS patients [19], and therefore may contribute to reduced oestrogen levels in humans with PCOS. Estrogens are not identified in our analysis of PCOS rodent models, potentially by Aα5 contributing to further inhibition of aromatase or by glucuronidation of androgens, or a combination of both. In our analysis, Aα5 is increased in LET-treated rats, but completely absent in DHT treated rats. Aα5 was absent in two biological replicates for the DHT-Control rat group and detected at very low levels in the LET-Control rats (**Figure 2C**). This may be due to LET-mediated inhibition of CYP19A mediating metabolic adaptations that are not mimicked by DHT administration [57].

Androgens are involved in the expression of the enzyme Cyclooxygenase-2 [58], which metabolises AA to prostaglandins. Abnormal Cyclooxygenase-2 levels are associated with infertility, failure of ovulation and disorders of oocyte implantation [59]. Androgens, γ-LA, an Ω-FA capable of producing AA, and AA itself are increased in certain PCOS subtypes [60]. In one rat model of PCOS, induced by alendronate, prostaglandins are increased in ovarian tissue [61]. These associations suggest a strong link between steroid hormones and mediators of inflammation in PCOS.

Leukotriene A_4_ (LTA_4_), produced from AA, can be metabolised to either Leukotriene B_4_ (LTB_4_) or LTC_4_. LTB_4_ is increased in PCOS patients [31]. In both, LET- and DHT-treated rats, LTC_4_ is significantly decreased. Reduced LTC_4_ levels may be due to the production of LTB_4_ from LTA_4_ instead of LTC_4_.

We have discovered a significant decrease in the concentration of D-sphingosine after DHT-treatment, which indicates that sphingolipid metabolism is affected. However, we have not yet researched other lipids in this metabolism pathway, in particular ceramides, which may be involved in this syndrome [32].

In conclusion, our method was able to simultaneously measure 22 lipids spanning the metabolic pathways of steroids, steroid hormones, bile acids, FA-based eicosanoid precursors, eicosanoids, and sphingolipids. We have quantified 12 of these metabolites, with significant changes in the amounts of LTC_4_, 25-hydroxycholesterol and 4-cholesten-3-one in the plasma of rodent models of PCOS. These lipids may provide alternative biomarkers for the diagnosis of this syndrome. However, the models show markedly different lipid profiles to each other. Therefore, this study proposes that caution is required when assessing the results from different models due to distinct lipid differences, as well as the presence of novel changes that are yet to be validated in humans.

### Data Availability Statement

Data is available upon request, please contact Hannah Scott at the University of Oxford at.

## Acknowledgments

Many thanks to Danielle Wellington (Chinese Academy of Medical Sciences) for contributing to the editing of the abstract and to James Dunford (Nuffield Department of Orthopaedics, Rheumatology and Musculoskeletal Sciences) for providing the standards from the Oppermann lab.

## Supplementary Materials

The supporting information can be downloaded at: xxxxxxxxxxxxxxxxx.

## Author Contributions

Conceptualisation and writing—original draft preparation, H.C.S., M.F., J.M., and B.M.K.; methodology, H.C.S, M.F., J.M., and G.B.; validation, formal analysis and visualisation, H.C.S., M.F., J.M.; software and data curation, H.C.S. and Z.Y.; investigation, H.C.S., M.F., J.M., D.O’B., C.B., and C.L.; resources, supervision, project administration and funding acquisition, B.M.K., M.F., M.P., U.O., and A.P.F.; writing—review and editing, H.C.S., D.O’B., A.P.F., and B.M.K. All authors have read and agreed to the published version of the manuscript.

## Abbreviations

γ-LA: γ-linolenic acid
3αAP/AP: 3α-Hydroxy-5α-Pregnane-20-one
3β-HSD: 3β-hydroxysteroid dehydrogenase
5-HPETE: 5-hydroperoxyeicosatetraenoic acid
A4: 4-Androstene-3,17-dione
Aα5: 5α-Androstane-3,17-dione
AA: Arachidonic acid
AKR1C1/2/4: 3α-hydroxysteroid dehydrogenase class 1/2/4
ALOX5: Arachidonate 5-lipoxygenase
ASAH1/2/ ACER1/2: neutral ceramidase
AUC: Area under the curve
CE: collisional energy
CH25H: Cholesterol 25-hydroxylase b
choD: cholesterol oxidase
COX-2: Cyclooxygenase 2
CV: coefficient of variation
CYP11: Steroid 11β-monooxygenase
CYP11A: Cytochrome P450 family 11 subfamily A
CYP11B1: Steroid 11β-monooxygenase
CYP142: cholest-4-en-3-one 26-monooxygenase
CYP17A: Steroid 17α-monooxygenase
CYP19A1: Cytochrome P450 aromatase
CYP21A: Steroid 21-monooxygenase
CYP2E1: Cytochrome P450 family 2 subfamily E1
DHCR7: 7-Dehydrocholesterol reductase
DHEA: Dehydroepiandrosterone
DHEA-S: Dehydroepiandrosterone-sulphate
DHT/5α-DHT: 5α-Dihydrotestosterone
DHT-C: Dihydrotestosterone control
diOHchol: Dihydrocholesterol
D-Sph/Sph: D-Sphingosine
dMRM: Dynamic multiple reaction monitoring
E1: Estrone
E1.1.1.51: 3 (or 17) β-Hydroxysteroid dehydrogenase
E2: β-Estradiol
E3: Estriol
E5.3.3.1: steroid Δ-isomerase
GABA_A_: γ-Aminobutyric acid type A
GALC: Galactosylceramidase
GBA: non-lysosomal glucosylceramidase
GGT1/5: leukotriene-C_4_ hydrolase
GPX4: phospholipid-hydroperoxide glutathione peroxidase
H_2_O: Water
HSD3B: 3β-Hydroxysteroid dehydrogenase
HSD11B1/2: Hydroxysteroid 11-β-dehydrogenase isozyme 1/2
HSD17B1/2: 17β-Estradiol 17-dehydrogenase 1/2
HSD17B3: Testosterone 17β-dehydrogenase (NADP+)
HSD17B11: Hydroxysteroid 17β-dehydrogenase 11
IM: ion mobility
IPA: Isopropanol
LC-dMRM-MS: Liquid chromatography-dynamic multiple reaction monitoring-mass spectrometry
LET: Letrozole
LET-C: Letrozole control
LLOD: Lower limit of detection
LLOQ: Lower limit of quantification
LTA_4_: Leukotriene A_4_
LTA_4_H: Leukotriene A_4_ hydrolase
LTB_4_: Leukotriene B_4_
LTC_4_: Leukotriene C_4_
LTC_4_S: Leukotriene C_4_ synthase
MAGL: Monoacylglycerol lipase
MeOH: Methanol
MRM: Multiple reaction monitoring
MTBE: Methyl tert-butyl ether
o.c.: on-column
PCOS: Polycystic ovarian syndrome
PLGA: poly-lactic-glycolic acid
PLPP1/2/3: Phosphatidate phosphatase 1/2/3
Preg: Pregnenolone
Prog: Progesterone
PTGR1/2: Prostaglandin reductase 1/2
PTGS1/2: prostaglandin-endoperoxide synthase 1
QC: Quality control
Qual: qualifier ion
Quant: Quantifier ion
RF: Response factor
S/N: Signal-to-noise ratio
SGMS1/2: sphingomyelin synthase
SGPP1/2: Sphingosine-1-phosphate phosphatase 1/2
SMPD1-4/ENPP7: ectonucleotide pyrophosphatase/phosphodiesterase family member 7
SPHK1/2: Sphingosine kinase 1/2
SRD5A1: 3-Oxo-5α-steroid 4-dehydrogenase 1
STD: standard
STS: Stearyl-sulfatase
SULT1E1: estrone sulfotransferase
SULT2B: Sulfotransferase family 2B member 1
TesI: 3-Oxo-5α-steroid 4-dehydrogenase
TIR: Transition Ion Ratios
UGT: uridine diphosphoglucuronosyltransferase
UGT8: Ceramide galactosyltransferase
ZADH2: Zinc binding alcohol dehydrogenase domain containing 2.

---

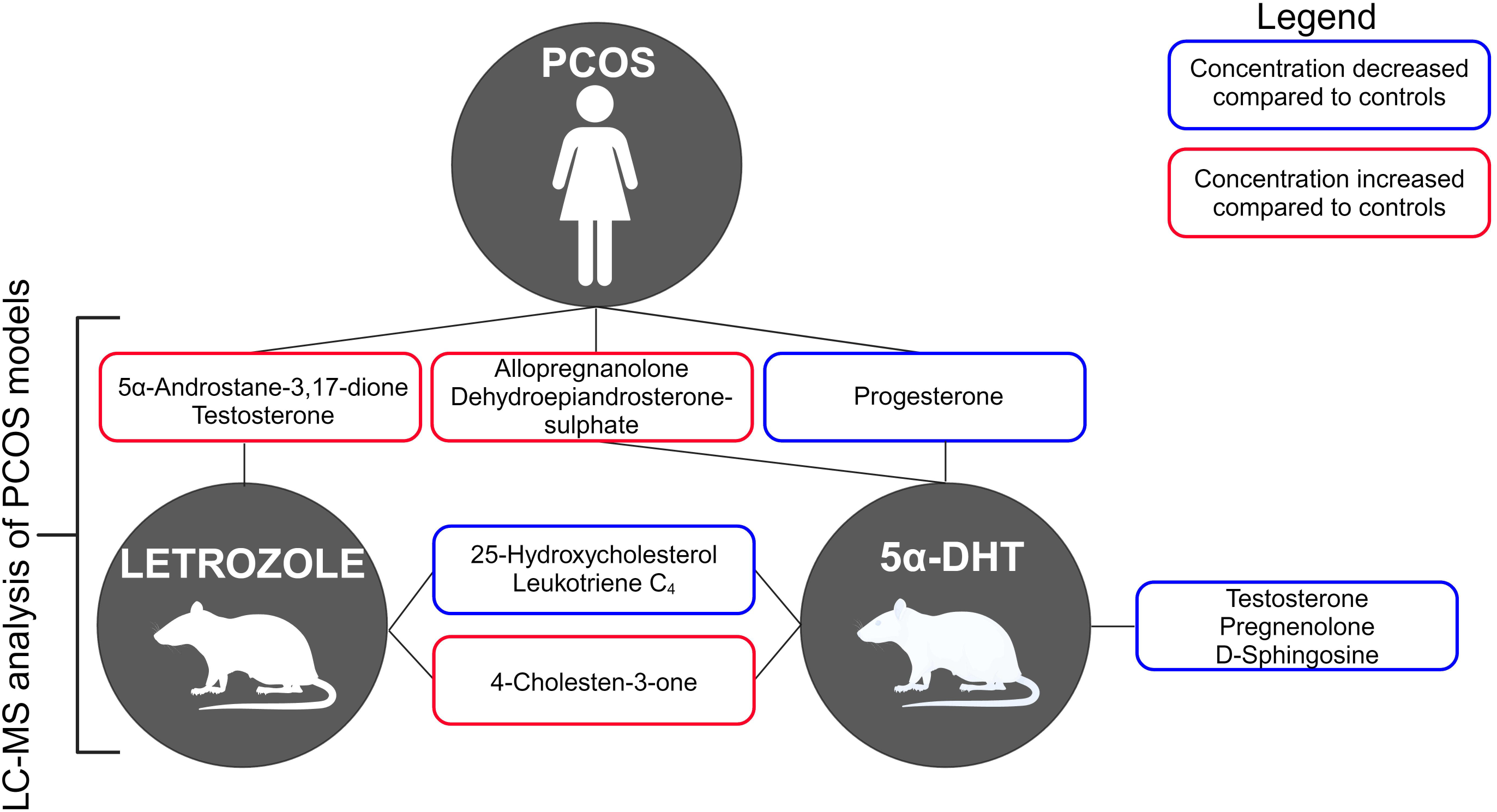

