## Supplementary figures and images for "Altered Hormone and Bioactive Lipid Plasma Profile in Rodent Models of Polycystic Ovarian Syndrome Revealed by Targeted Mass Spectrometry"

### Supplemental Figure 1

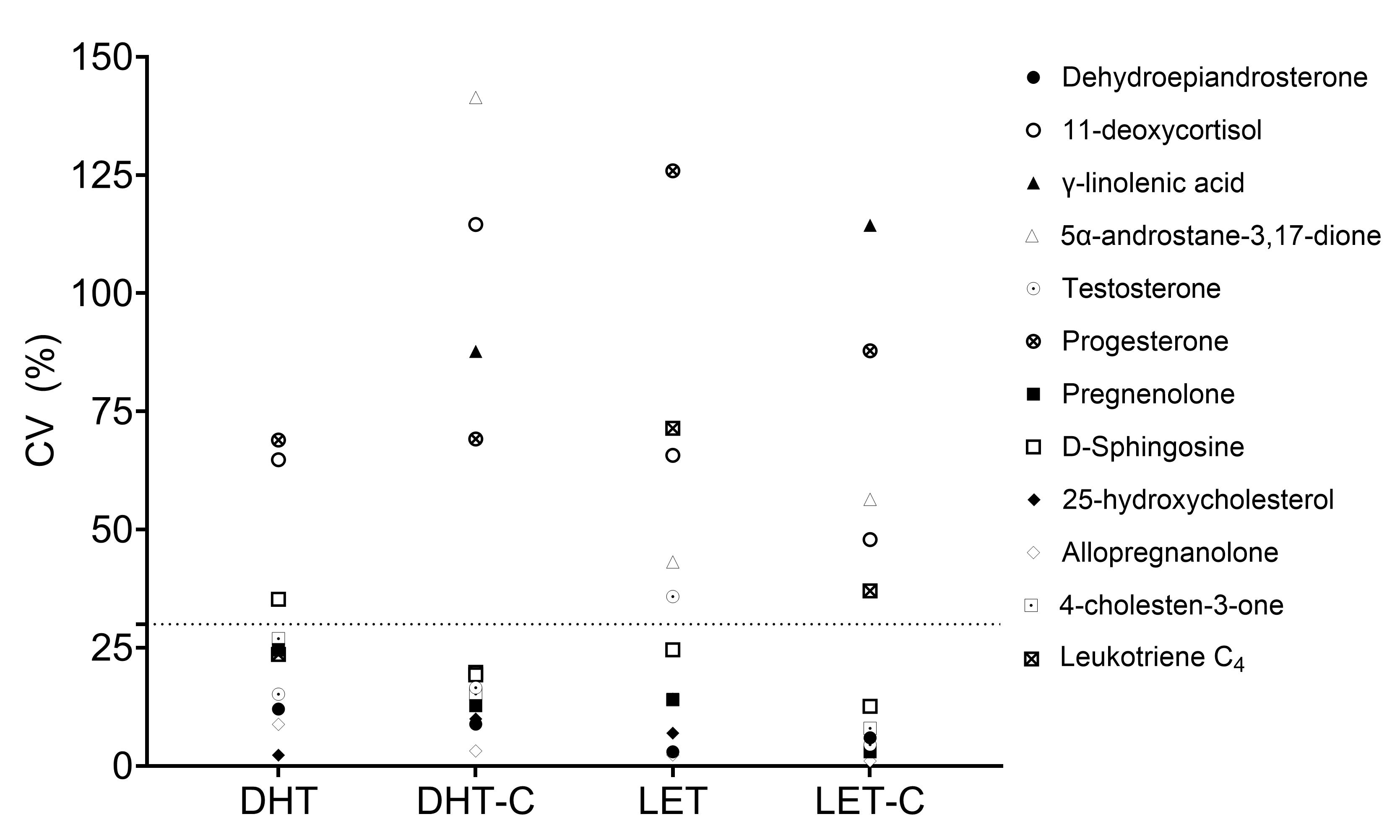

### Supplemental Figure 2

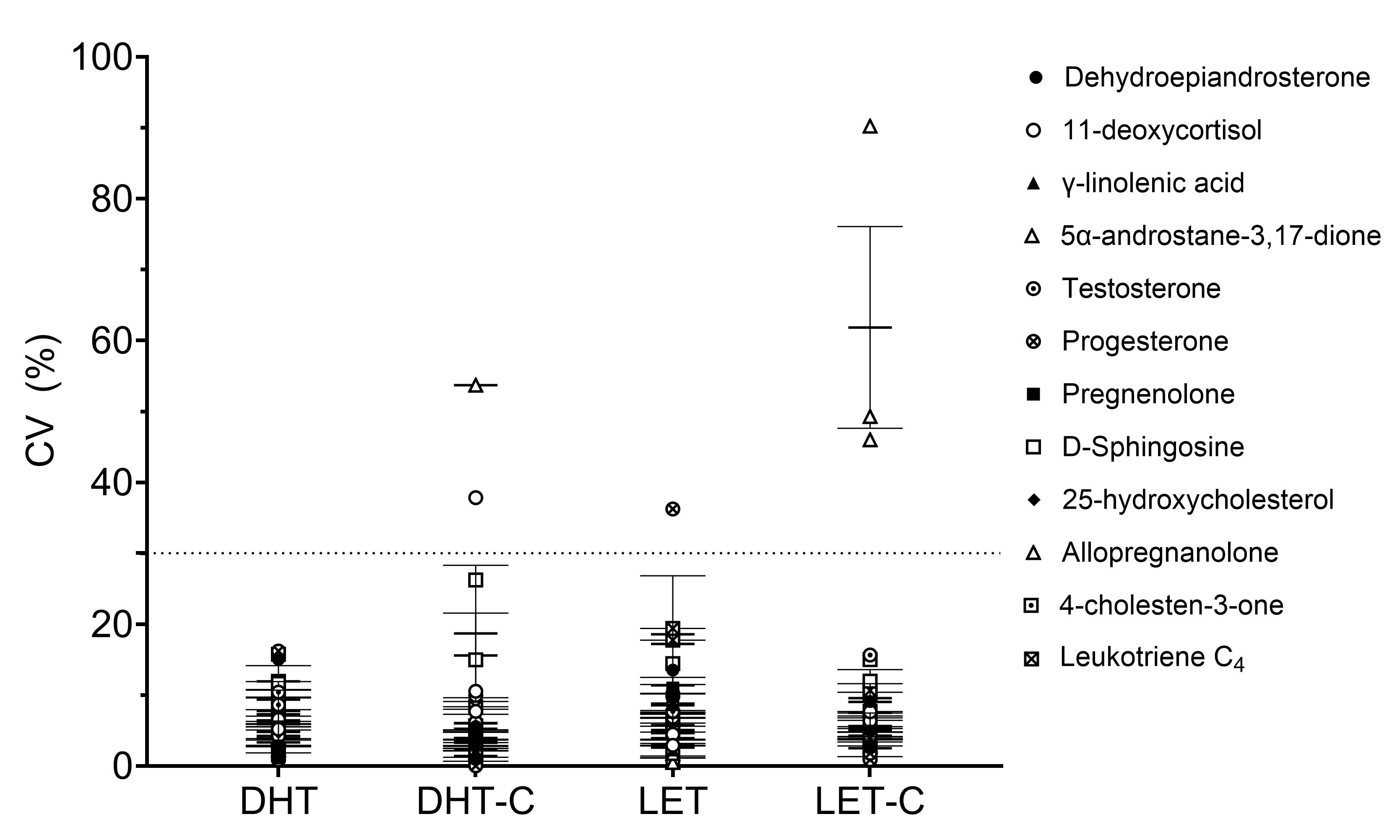

### Supplemental Figure 3

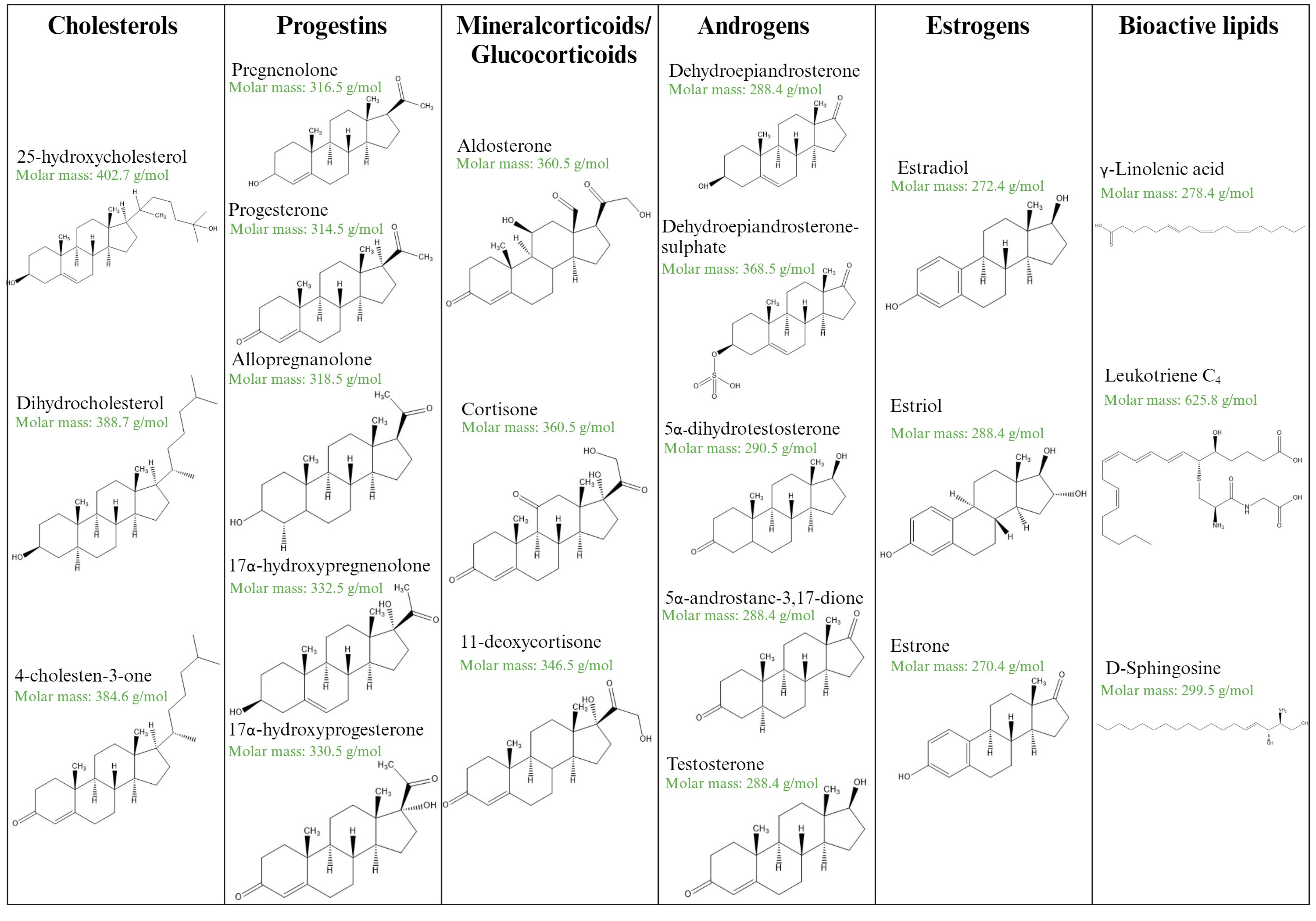

### Supplemental Figure 4A

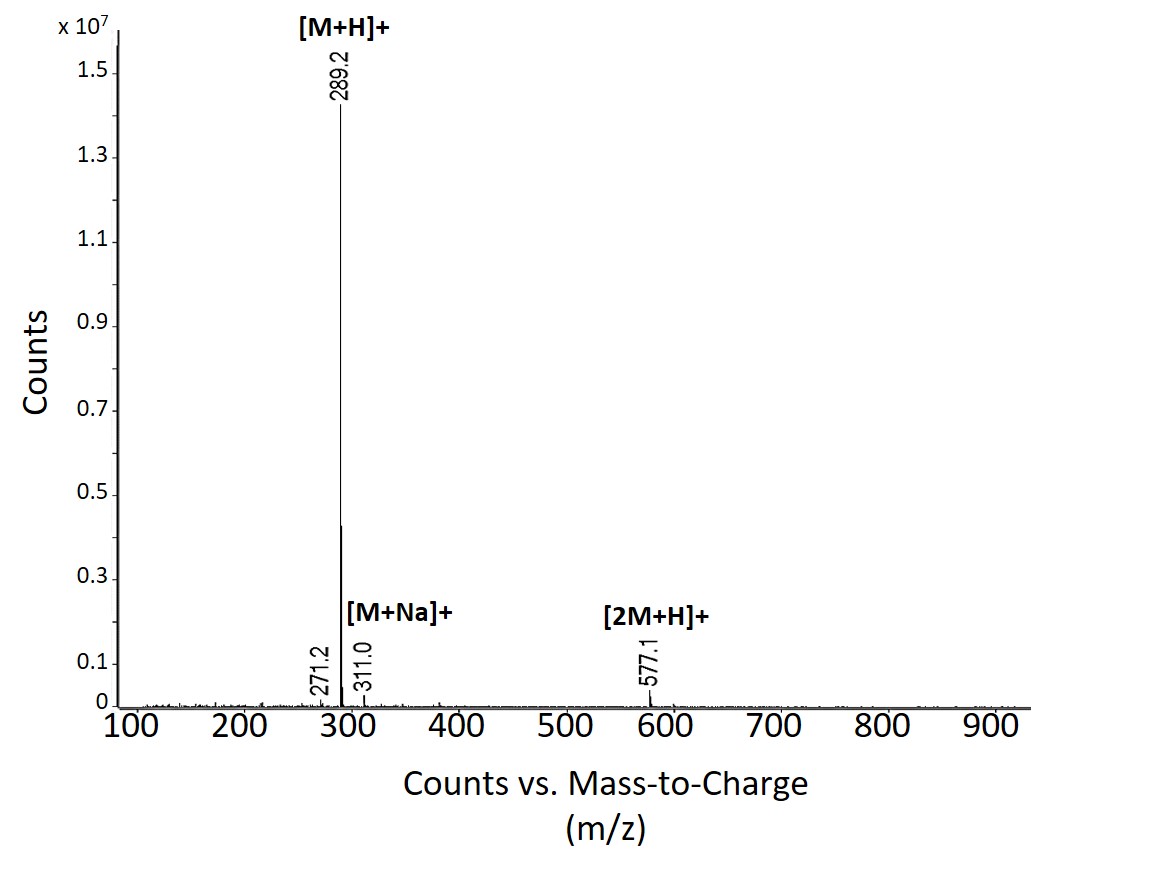

### Supplemental Figure 4B

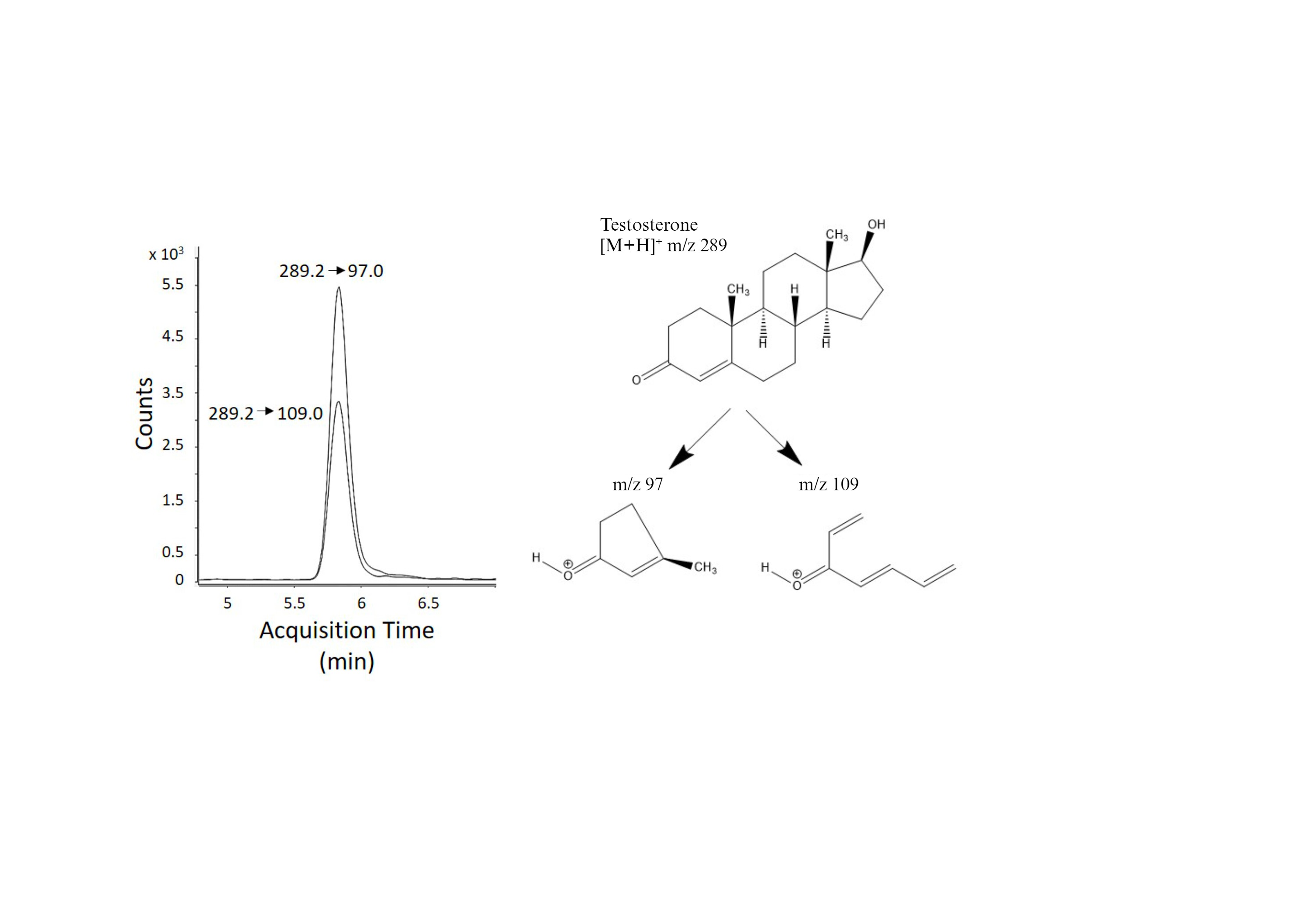

### Supplemental Figure 4C

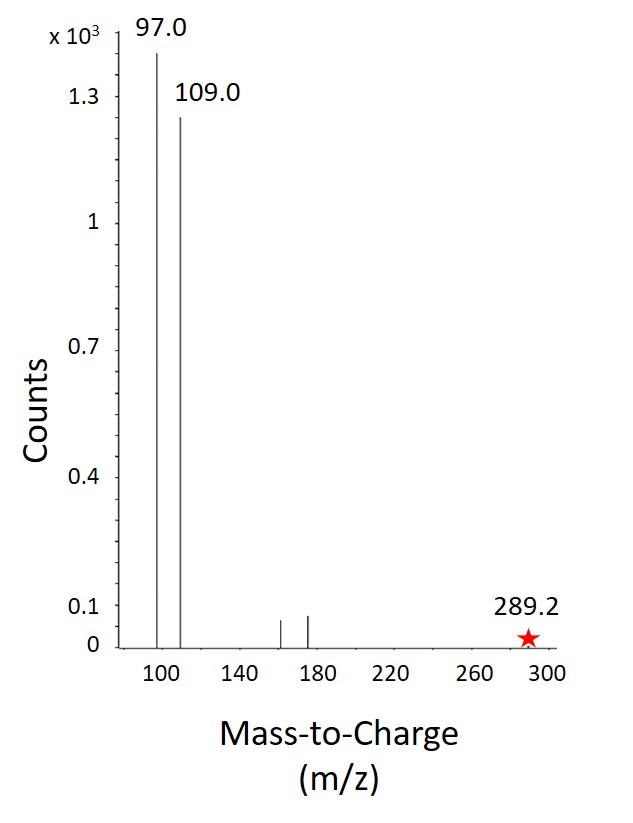

### Supplemental Figure 4D

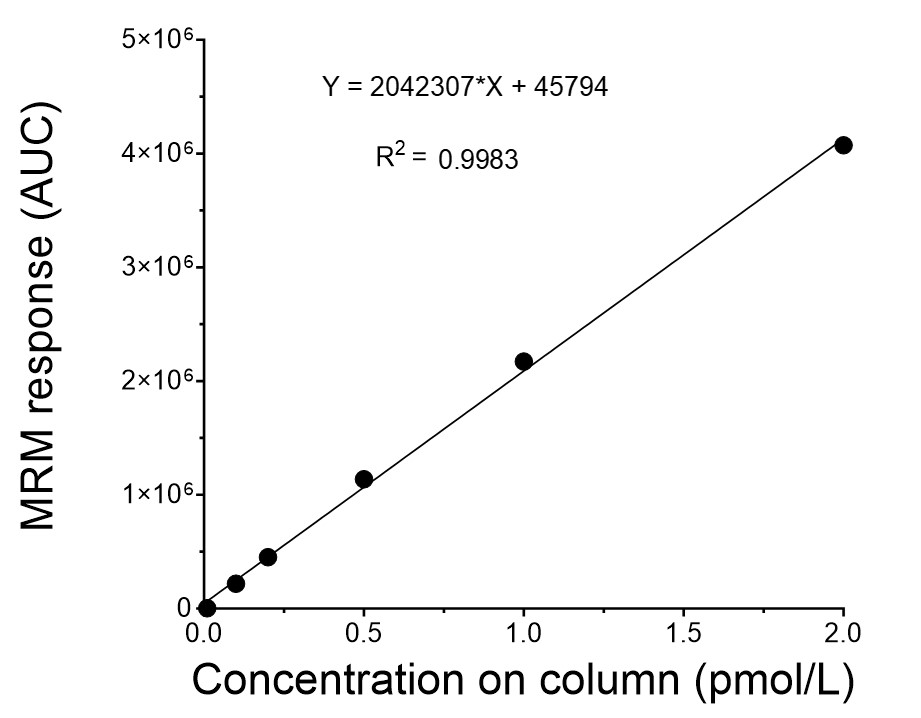

### Supplemental Figure 5

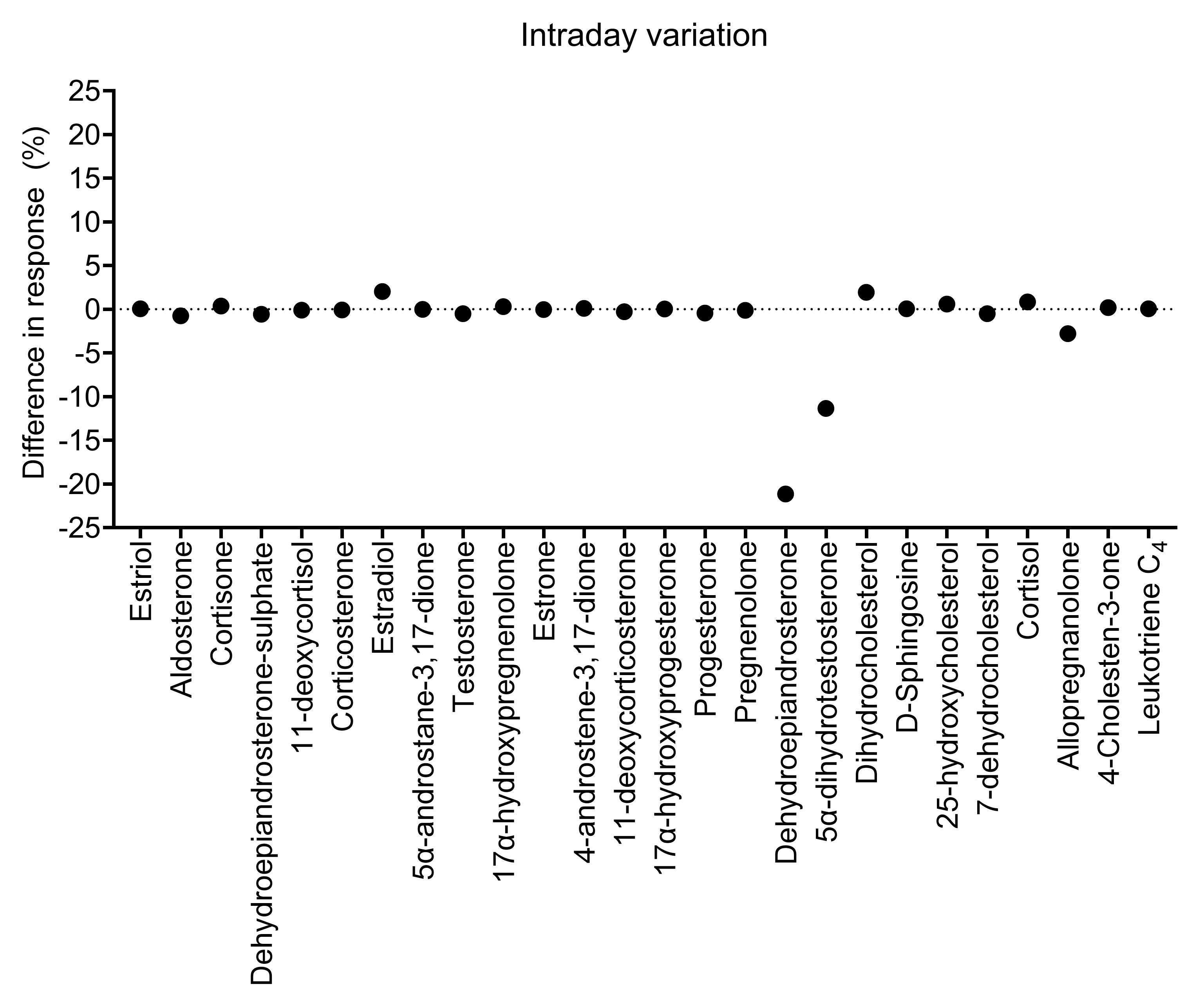
