## Supplementary material for "Altered Hormone and Bioactive Lipid Plasma Profile in Rodent Models of Polycystic Ovarian Syndrome Revealed by Targeted Mass Spectrometry": Legend Supplemental Figure 1

**Supplementary Figure 1**: ***Biological replicate analysis***. Samples are in biological triplicate (n=3). The coefficient of variation in percentage (CV %) is the standard deviation divided by the mean, multiplied by 100. The mean of the 3 biological replicates has been calculated for each lipid. LET-C represents non-treated control for Letrozole, LET is for Letrozole treated, DHT-C is a non-treated control for DHT, and DHT is Dihydrotestosterone treated. A CV of less than 30% represents a good biological reproducibility, less than 20% is very good reproducibility and less than 10% is excellent reproducibility.
