## Supplementary material for "Altered Hormone and Bioactive Lipid Plasma Profile in Rodent Models of Polycystic Ovarian Syndrome Revealed by Targeted Mass Spectrometry": Legend Supplemental Figure 2

**Supplementary Figure 2**: ***Technical replicate analysis***. Technical replicates (x3) analysis of the samples that are in biological triplicate (x3), n=9. The coefficient of variation in percentage (CV %) is the standard deviation divided by the mean, multiplied by 100. The mean of the 3 technical replicates, the mean of the means (black line) and standard error of the means (SEM, error bars) has been calculated for each lipid. LET-C represents non-treated control for Letrozole, LET is for Letrozole treated, DHT-C is a non-treated control for DHT, and DHT is Dihydrotestosterone treated. A CV of less than 30% represents a good technical reproducibility, less than 20% is very good reproducibility and less than 10% is excellent reproducibility.
