## Supplementary material for "Altered Hormone and Bioactive Lipid Plasma Profile in Rodent Models of Polycystic Ovarian Syndrome Revealed by Targeted Mass Spectrometry": Legend Supplemental Figure 3

**Supplementary Figure 3**: ***Chemical structures of the analysed lipids.*** Lipids analysed with this method include those within the groups of Cholesterols, Progestins, Glucocorticoids and Mineralocorticoids, Estrogens and Bioactive lipids. Their molecular mass is given in g/mol.
