## Supplementary material for "Altered Hormone and Bioactive Lipid Plasma Profile in Rodent Models of Polycystic Ovarian Syndrome Revealed by Targeted Mass Spectrometry": Legend Supplemental Figure 4A

**Supplementary Figure 4A**: ***The MS^2^ ion spectra of Testosterone (T).*** Testosterone has a molar mass of 288.431. In positive electrospray ionisation the protonated mass was identified at 289 m/z, the sodium adduct at 311 m/z and a protonated dimer at 577.1 m/z. There is a fragment ion caused by in-source fragmentation at 271.2 m/z. We used this information to cross reference our spectra with those found in databases (such as HMDB and LipidMaps) to corroborate identification of the correct lipid. We were also able to establish the retention time of the lipid in MS^2^ mode, to use in the setting of a retention time window for the dMRM transition.
