## Supplementary material for "Altered Hormone and Bioactive Lipid Plasma Profile in Rodent Models of Polycystic Ovarian Syndrome Revealed by Targeted Mass Spectrometry": Legend Supplemental Figure 4B

**Supplementary Figure 4B**: ***dMRM chromatogram of Testosterone (T).*** The protonated molecule at m/z 289.2 is exposed to a collisional energy 28 eV to obtain the fragment (product ion) of m/z 97.0, and collisional energy 32 eV for a product ion of 109.0 m/z. The product ion m/z 97.0 is the most abundant and is therefore used as the quantifier (quant) ion for determining concentration.
