## Supplementary material for "Altered Hormone and Bioactive Lipid Plasma Profile in Rodent Models of Polycystic Ovarian Syndrome Revealed by Targeted Mass Spectrometry": Legend Supplemental Figure 4C

**Supplementary Figure 4C*: dMRM spectra of Testosterone (T)***. The protonated ion (precursor ion) is indicated by the red star at 289.2 m/z. The fragments (product ions) are at *m/z* 97.0 and 109.0. The product ion 97.0 m/z is the most abundant and is therefore used as the quantifier (quant) ion for determining concentration. The use of ammonium acetate (CH_3_COONH_4_) as a mobile phase modifier can create [M+NH_4_]^+^ adducts, as seen in the analysis of 17α-hydroxypregnenolone (17αOHpreg) and 25-hydroxycholesterol (25OHchol), whereby some dMRM precursor ions are [M+NH_4_]^+^ adducts (a mass increase of 18).
