## Supplementary material for "Altered Hormone and Bioactive Lipid Plasma Profile in Rodent Models of Polycystic Ovarian Syndrome Revealed by Targeted Mass Spectrometry": Legend Supplemental Figure 4D

**Supplementary Figure 4D**: ***Response curve of Testosterone.*** Calibration curves (response curves) are generated using the response of the product ion in the mass spectrometer. On the X axis is the concentration, on the Y axis is the response (the counts), also known as the area under the curve (AUC). The trendline equation (Y=MX+C) is presented and accuracy of the trendline is shown by the R^2^.
