## Supplementary material for "Altered Hormone and Bioactive Lipid Plasma Profile in Rodent Models of Polycystic Ovarian Syndrome Revealed by Targeted Mass Spectrometry": Legend Supplemental Figure 5

**Supplementary Figure 5**: ***Method reproducibility (intraday analysis).*** Lipid standards were combined and analysed twice in technical triplicate at 100 fmol o.c. The mean percentage change from the first analysis was calculated (as %) for each standard. Most lipids had an intraday variation of less than +/- 5%.
