## Supplemental Table 1 for "Altered Hormone and Bioactive Lipid Plasma Profile in Rodent Models of Polycystic Ovarian Syndrome Revealed by Targeted Mass Spectrometry"

| **Lipid** | **Formula** | **Molar mass (g/mol)** | **Retention time (min)** | **Polarity** | **dMRM Precursor ion** | **Precursor ion type** | **dMRMProduct ions** | **Collisional energy (eV)** | **LLOD** | **LLOQ** |
| --- | --- | --- | --- | --- | --- | --- | --- | --- | --- | --- |
| Aldosterone | C_21_H_28_O_5_ | 360.45 | 1.95 | Positive | 361.45 | [M+H]+ | **91** | 44 | 100 fmol/L | 500 fmol/L |
|  |  |  |  |  | 361.5 | [M+H]+ | 79.2 | 72 | <150 ymol/L | 150 ymol/L |
| Estriol | C_18_H_24_O_3_ | 288.39 | 1.97 | Negative | 287 | [M-H]- | **171** | 40 | 100 pmol/L | 100 pmol/L |
|  |  |  |  |  | 287 | [M-H]- | 145 | 44 | 100 pmol/L | 150 pmol/L |
| Dehydroepiandrosterone-sulphate | C_19_H_28_O_5_S | 368.49 | 2.2 | Negative | 367 | [M-H]- | **97** | 40 | < 1.5 zmol/L | < 1.5 zmol/L |
| Cortisone | C_21_H_28_O_5_ | 360.45 | 2.45 | Positive | 361.5 | [M+H]+ | **163.1** | 28 | < 100 fmol/L | < 100 fmol/L |
|  |  |  |  |  | 361 | [M+H]+ | 91 | 64 | < 100 fmol/L | < 100 fmol/L |
|  |  |  |  |  | 361 | [M+H]+ | 163 | 24 | < 100 fmol/L | < 100 fmol/L |
| 11-deoxycortisol | C_21_H_30_O_4_ | 346.46 | 3.4 | Positive | 347.47 | [M+H]+ | **97.1** | 32 | 50 ymol/L | 50 fmol/L |
|  |  |  |  |  | 313 | Unknown | 283 | 30 | 50 pmol/L | 2 nmol/L |
|  |  |  |  |  | 313 | Unknown | 285 | 26 | 50 pmol/L | 2 nmol/L |
| γ-linolenic acid | C_18_H_30_O_2_ | 278.43 | 5.05 | Positive | 279.2 | [M+K]+ | 149.2 | 20 | < 100 zmol/L | < 100 zmol/L |
|  |  |  |  |  | 279.2 | [M+K]+ | 77.2 | 60 | 100 amol/L | 100 amol/L |
|  |  |  |  |  | 279.2 | [M+K]+ | **201.0** | 28 | < 100 zmol/L | < 100 zmol/L |
| 5α-androstane-3,17-dione | C_19_H_28_O_2_ | 288.43 | 5.1 | Positive | 327.2 | [M+K]+ | 41.2 | 76 | < 25 zmol/L | 25 zmol/L |
|  |  |  |  |  | 327.2 | [M+K]+ | 43.1 | 44 | < 1 ymol/L | > 1 ymol/L |
|  |  |  |  |  | 289.4 | [M+H]+ | 160.9 | 24 | < 1 ymol/L | > 1 ymol/L |
|  |  |  |  |  | 289.4 | [M+H]+ | **175** | 24 | < 1 ymol/L | > 1 ymol/L |
|  |  |  |  |  | 327.2 | [M+K]+ | 58.1 | 48 | < 1 ymol/L | > 1 ymol/L |
| Testosterone | C_19_H_28_O_2_ | 288.42 | 5.19 | Positive | 289.2 | [M+H]+ | **97** | 28 | 262.5 ymol/L | 87.5 zmol/L |
|  |  |  |  |  | 289 | [M+H]+ | 109 | 32 | 262.5 ymol/L | 87.5 zmol/L |
| Estradiol | C_18_H_24_O_2_ | 272.40 | 5.29 | Negative | 271 | [M-H]- | **145** | 48 | 1 amol/L | > 1 amol/L |
|  |  |  |  |  | 271 | [M-H]- | 183 | 52 | < 100 pmol/L | 100 pmol/L |
| Estrone | C_18_H_22_O_2_ | 270.37 | 6.68 | Negative | 269 | [M-H]- | **145** | 40 | < 1 pmol/L | 1 pmol/L |
|  |  |  |  |  | 269 | [M-H]- | 253 | 40 | 33.3 fmol/L | < 166.5 fmol/L |
| 17α-hydroxypregnenolone | C_21_H_32_O_3_ | 332.48 | 7.1 | Positive | 331 | Unknown | **287** | 19 | < 1 fmol/L | < 1 fmol/L |
|  |  |  |  |  | 350.5 | [M+NH_4_]+ | 45.1 | 44 | < 1 fmol/L | 1 fmol/L |
|  |  |  |  |  | 355.2 | [M+Na]+ | 73.1 | 36 | < 2 fmol/L | 2 fmol/L |
|  |  |  |  |  | 350.5 | [M+NH_4_]+ | 104.9 | 24 | 1 pmol/L | > 1 pmol/L |
|  |  |  |  |  | 350.5 | [M+NH_4_]+ | 186.9 | 16 | 1 pmol/L | > 1 pmol/L |
| 17α-hydroxyprogesterone | C_21_H_30_O_3_ | 330.46 | 7.23 | Positive | 331.23 | [M+H]+ | **97** | 32 | < 700 amol/L | 700 amol/L |
|  |  |  |  |  | 331 | [M+H]+ | 109 | 28 | 700 amol/L | 1.2 fmol/L |
| Progesterone | C_21_H_30_O_2_ | 314.47 | 11.5 | Positive | 315 | [M+H]+ | 109 | 40 | 500 amol/L | 750 amol/L |
|  |  |  |  |  | 315.2 | [M+H]+ | **97.1** | 24 | 250 amol/L | 500 amol/L |
| Pregnenolone | C_21_H_32_O_2_ | 316.49 | 13.1 | Positive | 299 | [M+H-H_2_O]+ | **281** | 20 | < 10 amol/L | 10 amol/L |
|  |  |  |  |  | 339.5 | [M+Na]+ | 43.2 | 48 | < 500 fmol/L | 500 fmol/L |
| 5α-dihydrotestosterone | C_19_H_30_O_2_ | 290.45 | 14.9 | Positive | 313.4 | [M+Na]+ | **59** | 32 | 10 ymol/L | > 10 ymol/L |
|  |  |  |  |  | 313.4 | [M+Na]+ | 105.1 | 16 | < 1 ymol/L | < 1 ymol/L |
|  |  |  |  |  | 313.4 | [M+Na]+ | 269.1 | 4 | < 1 ymol/L | < 1 ymol/L |
| Dihydrocholesterol | C_27_H_48_O | 388.67 | 16.3 | Positive | 411.7 | [M+Na]+ | **59.1** | 16 | < 50 ymol/L | < 50 ymol/L |
| D-sphingosine | C_18_H_37_NO_2_ | 299.50 | 16.7 | Positive | 300.5 | [M+H]+ | **282.3** | 8 | 500 ymol/L | 2 zmol/L |
|  |  |  |  |  | 300.5 | [M+H]+ | 55.1 | 40 | 5 fmol/L | 10 fmol/L |
|  |  |  |  |  | 300.5 | [M+H]+ | 56.1 | 32 | < 5 amol/L | 5 amol/L |
| Dehydroepiandrosterone | C_19_H_28_O_2_ | 288.42 | 18.1 | Positive | 311.4 | [M+Na]+ | **191.1** | 16 | < 1 ymol/L | 400 zmol/L |
| 25-hydroxycholesterol | C_27_H_46_O_2_ | 402.65 | 19.7 | Positive | 420.7 | [M+NH_4_]+ | **171.1** | 20 | < 1 ymol/L | < 1 ymol/L |
|  |  |  |  |  | 420.7 | [M+NH_4_]+ | 43.2 | 60 | < 1 ymol/L | < 1 ymol/L |
|  |  |  |  |  | 420.7 | [M+NH_4_]+ | 57 | 56 | < 1 ymol/L | < 1 ymol/L |
|  |  |  |  |  | 441.6 | [M+K]+ | 59 | 40 | < 1 ymol/L | < 1 ymol/L |
| Allopregnanolone | C_21_H_34_O_2_ | 318.50 | 23.5 | Positive | 341.5 | [M+Na]+ | **43.2** | 64 | < 50 ymol/L | < 50 ymol/L |
|  |  |  |  |  | 341.5 | [M+Na]+ | 41.1 | 76 | < 50 ymol/L | < 50 ymol/L |
|  |  |  |  |  | 341.5 | [M+Na]+ | 57 | 32 | < 50 ymol/L | < 50 ymol/L |
| 4-cholesten-3-one | C_27_H_44_O | 384.64 | 26.9 | Positive | 385.7 | [M+H]+ | **97** | 32 | < 50 ymol/L | < 50 ymol/L |
|  |  |  |  |  | 385.7 | [M+H]+ | 79 | 80 | < 50 ymol/L | < 50 ymol/L |
|  |  |  |  |  | 385.7 | [M+H]+ | 81 | 64 | 50 ymol/L | 100 ymol/L |
|  |  |  |  |  | 385.7 | [M+H]+ | 109.1 | 40 | < 50 ymol/L | < 50 ymol/L |
| Leukotriene C_4_ | C_30_H_47_N_3_O_9_S | 625.78 | 28.3 | Positive | 664.3 | [M+K]+ | **496.3** | 36 | < 50 ymol/L | < 50 ymol/L |
|  |  |  |  |  | 664.3 | [M+K]+ | 57.2 | 72 | < 50 ymol/L | < 50 ymol/L |
