## Supplementary material for "Altered Hormone and Bioactive Lipid Plasma Profile in Rodent Models of Polycystic Ovarian Syndrome Revealed by Targeted Mass Spectrometry": Legend Supplemental Table 1

**Supplementary Table 1**: ***Optimised transitions for LC-dMRM-MS.*** Lipid names, dMRM parameters and lower limits of quantification (LLOQ) are listed for all lipid standards analysed. The collisional energy used for the precursor-to-product ion transitions are indicated in electron volts (eV). The quantifier ion is indicated with bold text.
