## Supplemental Table 2 for "Altered Hormone and Bioactive Lipid Plasma Profile in Rodent Models of Polycystic Ovarian Syndrome Revealed by Targeted Mass Spectrometry"

| **Chromatography setting** | **Parameter** |
| --- | --- |
| Mobile phase composition | A = 40% ACN, 5mM Ammonium Acetate  B = 90% IPA, 10% ACN, 5mM Ammonium Acetate |
| Mobile phase flow rate (ml/min) | 0.2 |
| C18 column temperature (°C) | 40 |
| **Mass spectrometer source setting** | **Parameter** |
| Gas temperature (°C) | 280 |
| Gas flow (L/min) | 14 |
| Nebulizer (psi) | 20 |
| Sheath gas temperature (°C) | 250 |
| Sheath gas flow (L/min) | 11 |
| Capillary (V) | 4000 |
| Nozzle (V) | 1000 |
| **Mass spectrometer iFunnel setting** | **Parameter** |
| High pressure RF (V) | 150 |
| Low pressure RF (V) | 90 |
