## Supplementary material for "Altered Hormone and Bioactive Lipid Plasma Profile in Rodent Models of Polycystic Ovarian Syndrome Revealed by Targeted Mass Spectrometry": Legend Supplemental Table 2

**Supplementary Table 2**: ***Chromatography and mass spectrometry parameters.*** This table represents the instrumentation settings used in the analysis method and parameters applied. Settings are selected for the most favourable separation and ionisation of steroids, steroid hormones, eicosanoids and sphingolipids.
