## Supplemental Table 3 for "Altered Hormone and Bioactive Lipid Plasma Profile in Rodent Models of Polycystic Ovarian Syndrome Revealed by Targeted Mass Spectrometry"

| **Biological**  **replicate variation (CV)** | **DHEA-S** | **11DOC** | **LA** | **A5** | **TESTOS** | **PROGES** | **PREGNE** | **SPH** | **25OHCH** | **AP** | **4CHOL3** | **LTC4** |
| --- | --- | --- | --- | --- | --- | --- | --- | --- | --- | --- | --- | --- |
| **LET** | 2.98 | 65.70 | 14.18 | 43.13 | 35.85 | 125.92 | 14.08 | 24.59 | 6.92 | 2.34 | 14.00 | 71.43 |
| **LET-C** | 5.95 | 47.91 | 114.37 | 56.42 | 4.48 | 87.83 | 2.99 | 12.60 | 5.70 | 1.06 | 7.97 | 37.04 |
| **DHT** | 12.02 | 64.79 | 24.52 | ND | 15.16 | 68.97 | 24.62 | 35.26 | 2.28 | 8.78 | 26.95 | 23.60 |
| **DHT-C** | 8.84 | 114.57 | 87.69 | 141.42 | 16.56 | 69.18 | 12.76 | 19.25 | 9.93 | 3.17 | 15.22 | 19.78 |

*^DHEA-S: Dehydroepiandrosterone-sulphate, 11DOC: 11-deoxycortisol, LA: γ-linolenic acid, A5: 5α-androstane-3,17-dione, TESTOS: Testosterone, PROGES: Progesterone, PREGNE: Pregnenolone, SPH: D-sphingosine, 25OHCH: 25-hydroxycholesterol, AP: Allopregnanolone, 4CHOL3: 4-cholesten-3-one, LTC4: Leukotriene C4, ND: not detected^*
