## Supplementary material for "Altered Hormone and Bioactive Lipid Plasma Profile in Rodent Models of Polycystic Ovarian Syndrome Revealed by Targeted Mass Spectrometry": Legend Supplemental Table 3

**Supplementary Table 3**: ***Biological variation analysis***. Coefficient of variation of biological replicates: Samples are in biological triplicate. The mean and standard deviation (std.dev/SD) has been calculated for biological replicates, and the coefficient of variation in percentage (CV %), given, is the standard deviation divided by the mean, multiplied by 100. LET-Control represents non-treated control for Letrozole, LET is for Letrozole treated, DHT-Control is a non-treated control for DHT, DHT; Dihydrotestosterone treated.
