## Supplemental Table 4 for "Altered Hormone and Bioactive Lipid Plasma Profile in Rodent Models of Polycystic Ovarian Syndrome Revealed by Targeted Mass Spectrometry"

| **Technical replicate variation (CV)** | **Technical**  **replicate** | **DHEA-S** | **11DOC** | **LA** | **A5** | **TESTOS** | **PROGES** | **PREGNE** | **SPH** | **25OHCH** | **AP** | **4CHOL3** | **LTC4** |
| --- | --- | --- | --- | --- | --- | --- | --- | --- | --- | --- | --- | --- | --- |
| **LET** | 1 | 9.72 | 4.49 | 8.73 | 0.49 | 0.63 | 36.23 | 9.07 | 5.70 | 1.51 | 0.63 | 2.03 | ND |
|  | 2 | 13.52 | 2.91 | 2.26 | 1.76 | 6.86 | 5.17 | 6.32 | 14.43 | 1.32 | 1.76 | 4.81 | 17.76 |
|  | 3 | 10.86 | 9.81 | 3.63 | 9.49 | 7.69 | 10.22 | 10.97 | 5.53 | 4.94 | 6.52 | 6.99 | 19.41 |
| **LET-C** | 1 | 4.96 | 6.36 | 2.85 | 46.01 | 15.66 | 4.81 | 5.39 | 14.99 | 4.73 | 5.71 | 3.90 | 10.12 |
|  | 2 | 9.08 | 2.70 | 6.75 | 49.28 | 2.69 | 0.88 | 5.13 | 1.73 | 3.26 | 2.72 | 7.70 | 10.65 |
|  | 3 | 2.89 | 7.63 | 6.80 | 90.23 | 4.14 | 1.78 | 4.57 | 11.95 | 4.85 | 2.94 | 3.83 | 6.31 |
| **DHT** | 1 | 15.04 | 6.58 | 5.40 | ND | 4.25 | 16.20 | 5.29 | 9.67 | 4.92 | 4.03 | 5.39 | 11.92 |
|  | 2 | 2.65 | 2.64 | 6.65 | ND | 8.62 | 0.83 | 6.98 | 11.64 | 6.08 | 3.63 | 4.68 | 15.77 |
|  | 3 | 1.10 | 5.19 | 5.64 | ND | 10.43 | 4.86 | 7.03 | 6.77 | 1.70 | 2.28 | 1.62 | 8.10 |
| **DHT-C** | 1 | 4.00 | 7.70 | 10.73 | ND | 9.95 | 0.02 | 4.56 | 26.23 | 4.00 | 4.06 | 4.23 | 8.36 |
|  | 2 | 5.54 | 10.53 | 2.97 | ND | 6.16 | 1.60 | 1.22 | 5.57 | 3.06 | 5.89 | 1.80 | 4.02 |
|  | 3 | 0.89 | 37.85 | 2.06 | 53.72 | 2.05 | 2.61 | 5.45 | 14.97 | 2.71 | 1.85 | 0.77 | 5.65 |

*^DHEA-S: Dehydroepiandrosterone-sulphate, 11DOC: 11-deoxycortisol, LA: γ-linolenic acid, A5: 5α-androstane-3,17-dione, TESTOS: Testosterone, PROGES: Progesterone, PREGNE: Pregnenolone, SPH: D-sphingosine, 25OHCH: 25-hydroxycholesterol, AP: Allopregnanolone, 4CHOL3: 4-cholesten-3-one, LTC4: Leukotriene C4, ND: not detected^*
