## Supplementary material for "Altered Hormone and Bioactive Lipid Plasma Profile in Rodent Models of Polycystic Ovarian Syndrome Revealed by Targeted Mass Spectrometry": Legend Supplemental Table 4

**Supplementary Table 4**: ***Technical variation analysis.*** Coefficient of variation in technical replicates: Samples are in biological triplicate. The mean and standard deviation (std.dev) has been calculated for biological replicates, the coefficient of variation in percentage (CV %), given, is the standard deviation divided by the mean, multiplied by 100. LET-Control represents non-treated control for letrozole, LET is for Letrozole treated, DHT-Control is a non-treated control for DHT, DHT; Dihydrotestosterone treated.
