## Supplemental Table 5 for "Altered Hormone and Bioactive Lipid Plasma Profile in Rodent Models of Polycystic Ovarian Syndrome Revealed by Targeted Mass Spectrometry"

| **Lipid in standard mixture** | **Equation** | **R^2^** |
| --- | --- | --- |
| 11-Deoxycortisol | Y = 7280173*X + 156768 | 0.999 |
| 17α-hydroxypregnenolone | Y = 9314*X + 40.90 | 0.996 |
| 17α-hydroxyprogesterone | Y = 1362106*X - 33303 | 0.999 |
| 17-β-Estradiol | Y = 1763*X - 71.64 | 0.989 |
| 25-Hydroxycholesterol | Y = 3504232*X + 2865777 | 0.958 |
| 3α-Hydroxy-5α-Pregnan-20-one | Y = 4005*X + 1045 | 0.928 |
| 4-Cholesten-3-one | Y = 10280*X + 3683 | 0.921 |
| 5α-Androstane-3,17-dione | Y = 5086188*X + 13211 | 0.999 |
| 5α-Dihydrotestosterone | Y = 122.6*X + 64.79 | 0.917 |
| Aldosterone | Y = 76127*X + 6027 | 0.970 |
| Cortisone | Y = 917395*X + 21821 | 0.998 |
| Dehydroepiandrosterone | Y = 508028*X + 87275 | 0.987 |
| Dehydroepiandrosterone-sulphate | Y = 55919*X - 1418 | 0.998 |
| Dihydrocholesterol | Y = 12392*X - 133.3 | 0.989 |
| D-Sphingosine | Y = 75622*X - 3191 | 0.996 |
| Estriol | Y = 1245*X + 60.14 | 0.977 |
| Estrone | Y = 4669*X - 83.46 | 0.999 |
| Leukotriene C_4_ | Y = 3131920*X + 7990556 | 0.922 |
| Pregnenolone | Y = 330778*X - 6979 | 0.998 |
| Progesterone | Y = 8054792*X - 88749 | 0.999 |
| Testosterone | Y = 5145893*X - 99878 | 0.999 |
| γ-Linolenic acid | Y = 11129427*X + 138911 | 0.998 |
