## Supplementary material for "Altered Hormone and Bioactive Lipid Plasma Profile in Rodent Models of Polycystic Ovarian Syndrome Revealed by Targeted Mass Spectrometry": Legend Supplemental Table 5

**Supplementary Table 5**: ***Response (calibration) curve equations and accuracy.*** Increasing concentrations of the standard lipid mix were injected and the response of the quantitation (quant) ion was monitored by observing the area under the curve (AUC). The MRM AUC’s were plotted against the on-column concentration to give response curves (calibration curves). Each response curve was plotted with a linear trendline. The values for the slope and intercept are given in the table above. R^2^; The accuracy of the fit of the linear trendline.
