## Supplemental Table 6 for "Altered Hormone and Bioactive Lipid Plasma Profile in Rodent Models of Polycystic Ovarian Syndrome Revealed by Targeted Mass Spectrometry"

| **Lipid** | **Formula** | **Precursor ion** | **Product ions (qualifiers)** | **Comments** | **Average ratio** | **SD** | **CV (%)** | **Isomer presence** |
| --- | --- | --- | --- | --- | --- | --- | --- | --- |
| Aldosterone | C_21_H_28_O_5_ | 361.5 | 79.2 | Not identified in any samples | NA | NA | NA | NA |
| Estriol | C_18_H_24_O_3_ | 287 | 145 | Not identified in any samples | NA | NA | NA | NA |
| Dehydroepiandrosterone-sulphate | C_19_H_28_O_5_S | NA | NA | Only quantifier transition | NA | NA | NA | NA |
| Cortisone | C_21_H_28_O_5_ | 361 | 91 | Not identified in any samples | NA | NA | NA | NA |
|  |  | 361 | 163 | Not identified in any samples | NA | NA | NA |  |
| 11-deoxycortisol | C_21_H_30_O_4_ | 313 | 283 | Not present in all analyses | NA | NA | NA | NA |
|  |  | 313 | 285 | Not present in all analyses | NA | NA | NA |  |
| γ-linolenic acid | C_18_H_30_O_2_ | 279.2 | 149.2 | Not present in all analyses | NA | NA | NA | NA |
|  |  | 279.2 | 77.2 | Not present in all analyses | NA | NA | NA |  |
| 5α-androstane-3,17-dione | C_19_H_28_O_2_ | 327.2 | 41.2 | Not present in all analyses | NA | NA | NA | NA |
|  |  | 327.2 | 43.1 | Not present in all analyses | NA | NA | NA |  |
|  |  | 327.2 | 58.1 | Not present in all analyses | NA | NA | NA |  |
|  |  | 289.4 | 160.9 | Not present in all analyses | NA | NA | NA |  |
| Testosterone | C_19_H_28_O_2_ | 289 | 109 | Not present in all analyses | 74.23 | 30.52 | 4.10 | Potentially but not DHEA as these are chromatographically separated |
| Estradiol | C_18_H_24_O_2_ | 271 | 183 | Not identified in any samples | NA | NA | NA | NA |
| Estrone | C_18_H_22_O_2_ | 269 | 253 | Not identified in any samples | NA | NA | NA | NA |
| 17α-hydroxypregnenolone | C_21_H_32_O_3_ | 350.5 | 45.1 | Not present in all analyses | NA | NA | NA | NA |
|  |  | 350.5 | 73.1 | Not present in all analyses | NA | NA | NA |  |
|  |  | 350.5 | 104.9 | Not present in all analyses | NA | NA | NA |  |
|  |  | 350.5 | 186.9 | Not present in all analyses | NA | NA | NA |  |
| 17α-hydroxyprogesterone | C_21_H_30_O_3_ | 331 | 109 | Not identified in any samples | NA | NA | NA | Potentially, could be 11-deoxycorticosterone |
| Progesterone | C_21_H_30_O_2_ | 315.2 | 109 | TIR analysis | 48.54 | 6.66 | 13.71 | Potentially |
| Pregnenolone | C_21_H_32_O_2_ | 339.5 | 43.2 | Not present in all analyses | NA | NA | NA | NA |
| 5α-dihydrotestosterone | C_19_H_30_O_2_ | 313.4 | 59 | Not present in all analyses | NA | NA | NA | Potentially, could be androsterone |
|  |  | 313.4 | 105.1 | Not present in all analyses | NA | NA | NA |  |
|  |  | 313.4 | 269.1 | Not present in all analyses | NA | NA | NA |  |
| Dihydrocholesterol | C_27_H_48_O | NA | NA | Only quantifier transition | NA | NA | NA | NA |
| D-sphingosine | C_18_H_37_NO_2_ | 300.5 | 55.1 | Not present in all analyses | NA | NA | NA | NA |
|  |  | 300.5 | 56.1 | Not present in all analyses | NA | NA | NA |  |
| Dehydroepiandrosterone | C_19_H_28_O_2_ | NA | NA | Only quantifier transition | NA | NA | NA | NA |
| 25-hydroxycholesterol | C_27_H_46_O_2_ | 420.7 | 43.2 | TIR analysis | 4.92 | 0.22 | 4.44 | No |
|  |  | 420.7 | 57 | TIR analysis | 36.29 | 1.44 | 3.96 |  |
|  |  | 441.6 | 59 | Not present in all analyses | NA | NA | NA |  |
| Allopregnanolone | C_21_H_34_O_2_ | 341.5 | 57 | Not present in all analyses | NA | NA | NA | Potentially |
|  |  | 341.5 | 41.1 | TIR analysis | 57.16 | 17.37 | 30.4 |  |
| 4-cholesten-3-one | C_27_H_44_O | 385.7 | 79 | Not present in all analyses | NA | NA | NA | No |
|  |  | 385.7 | 81 | Not present in all analyses | NA | NA | NA |  |
|  |  | 385.7 | 109.1 | TIR analysis | 84.81 | 4.99 | 5.89 |  |
| Leukotriene C_4_ | C_30_H_47_N_3_O_9_S | 664.3 | 57.2 | Not identified in any samples | NA | NA | NA | No |

*^NA = not applicable^*
