## Supplementary material for "Altered Hormone and Bioactive Lipid Plasma Profile in Rodent Models of Polycystic Ovarian Syndrome Revealed by Targeted Mass Spectrometry": Legend Supplemental Table 6

**Supplementary Table 6**: ***Transition ion ratio analysis of lipids identified in samples to identify potential isomers.*** TIR analysis requires the presence of both quantifier (qual) and qualifier (qual) ion transitions. A high average ratio of quant to qual ions indicates the presence of isomeric species. The reproducibility of the TIR average ratio is represented by the standard deviation (SD) and the coefficient of variation (CE). When the TIR analysis is not relevant or cannot be applied then it is represented by NA (not applicable).
