## Supplemental Table 7 for "Altered Hormone and Bioactive Lipid Plasma Profile in Rodent Models of Polycystic Ovarian Syndrome Revealed by Targeted Mass Spectrometry"

|  | **LET** | | | **LET-C** | | | **DHT** | | | **DHT-C** | | |
| --- | --- | --- | --- | --- | --- | --- | --- | --- | --- | --- | --- | --- |
|  | **MEAN** | **SD** | **N** | **MEAN** | **SD** | **N** | **MEAN** | **SD** | **N** | **MEAN** | **SD** | **N** |
| **DHEA-S** | 1.512 pmol/L | 0.19 | 9 | 1.46 pmol/L | 0.14 | 9 | 1.55 pmol/L | 0.24 | 9 | 1.33 pmol/L | 0.14 | 9 |
| **TESTOS** | 173.40 fmol/L | 67.31 | 9 | 22.69 fmol/L | 2.42 | 9 | 10.78 fmol/L | 1.94 | 9 | 15.33 fmol/L | 2.86 | 9 |
| **A5** | 140.13 fmol/L | 64.81 | 9 | 4.38 fmol/L | 4.22 | 9 | 190.76 fmol/L | 19.09 | 9 | 166.09 fmol/L | 9.49 | 9 |
| **PROGES** | 0.40 pmol/L | 0.54 | 9 | 0.58 pmol/L | 0.54 | 9 | 78.22 fmol/L | 57.21 | 9 | 410.56 fmol/L | 301.32 | 9 |
| **PREGNE** | 23.15 pmol/L | 4.17 | 9 | 22.07 pmol/L | 1.37 | 9 | 30.59 pmol/L | 8.27 | 9 | 43.49 pmol/L | 6.18 | 9 |
| **AP** | 246.21 pmol/L | 12.11 | 9 | 237.87 pmol/L | 10.58 | 9 | 190.76 pmol/L | 19.09 | 9 | 166.09 pmol/L | 9.49 | 9 |
| **25OHCH** | 15.71 pmol/L | 1.25 | 9 | 19.92 pmol/L | 1.52 | 9 | 15.37 pmol/L | 0.84 | 9 | 21.57 pmol/L | 2.39 | 9 |
| **11DOC** | 1.59 pmol/L | 1.12 | 9 | 1.11 pmol/L | 0.57 | 9 | 0.86 pmol/L | 0.60 | 9 | 0.44 pmol/L | 0.54 | 9 |
| **4CHOL3** | 385.20 pmol/L | 61.17 | 9 | 312.67 pmol/L | 32.62 | 9 | 678.30 pmol/L | 195.67 | 9 | 227.78 pmol/L | 37.55 | 9 |
| **LA** | 55.51 fmol/L | 9.10 | 9 | 224.56 fmol/L | 272.62 | 9 | 221.49 fmol/L | 59.48 | 9 | 201.72 fmol/L | 187.84 | 9 |
| **LTC4** | 1.51 pmol/L | 1.20 | 9 | 3.85 pmol/L | 1.57 | 9 | 6.09 pmol/L | 1.74 | 9 | 10.41 pmol/L | 2.28 | 9 |
| **SPH** | 2.11 pmol/L | 0.58 | 9 | 1.86 pmol/L | 0.33 | 9 | 1.61 pmol/L | 0.63 | 9 | 2.23 pmol/L | 0.58 | 9 |

*^DHEA-S: Dehydroepiandrosterone-sulphate, 11DOC: 11-deoxycortisol, LA: γ-linolenic acid, A5: 5α-androstane-3,17-dione, TESTOS: Testosterone, PROGES: Progesterone, PREGNE: Pregnenolone, SPH: D-sphingosine, 25OHCH: 25-hydroxycholesterol, AP: Allopregnanolone, 4CHOL3: 4-cholesten-3-one, LTC4: Leukotriene C4, N: number of analyses^*
