## Supplementary material for "Altered Hormone and Bioactive Lipid Plasma Profile in Rodent Models of Polycystic Ovarian Syndrome Revealed by Targeted Mass Spectrometry": Legend Supplemental Table 7

**Supplementary Table 7**: ***LET and DHT treatment of rats resulted in marked changes to the lipids profiled in rat plasma.*** The concentration (mean) of all analyses inclusive of technical triplicate of biological triplicates, the standard deviation (SD) and the number of analyses (N). The concentration units are given in each mean cell.
